# Signaling input from divergent pathways subverts malignant B-cell transformation

**DOI:** 10.1101/2020.03.12.989749

**Authors:** Lai N. Chan, Mark A. Murakami, Mark E. Robinson, Rebecca Caeser, Teresa Sadras, Jaewoong Lee, Kadriye Nehir Cosgun, Kohei Kume, Vishal Khairnar, Gang Xiao, Mohamed Ahmed, Eamon Aghania, Gauri Deb, Christian Hurtz, Seyedmehdi Shojaee, Chao Hong, Petri Pölönen, Matthew A. Nix, Zhengshan Chen, Chun Wei Chen, Jianjun Chen, Andreas Vogt, Merja Heinäniemi, Olli Lohi, Arun P. Wiita, Shai Izraeli, Huimin Geng, David M. Weinstock, Markus Müschen

**Author notes:** For correspondence: Markus Müschen, MD-PhD.

## Abstract

Malignant transformation typically involves multiple genetic lesions whose combined activity gives rise to cancer^1^. Our analysis of 1,148 patient-derived B-cell leukemia (B-ALL) samples revealed that individual mutations did not promote leukemogenesis unless they converged on one single oncogenic pathway characteristic for the differentiation status of these transformed B cells. Specifically, we show here the JAK/STAT5 signaling pathway supports the developmental stage-specific expansion of pro-B ALL whereas the ERK-pathway that of pre-B ALL. Mutations that were not aligned with the central oncogenic driver would activate divergent pathways and subvert malignant transformation. Oncogenic lesions in B-ALL frequently mimic survival and proliferation signals downstream of cytokine receptors (through activation of STAT5)^2-7^ or the pre-B cell receptor (through activation of ERK)^8-13^. STAT5- (372 cases) and ERK- (386 cases) activating lesions were frequently found but only co-occurred in ∼3% (37) of cases (*P*=2.2E-16). Single-cell mutation and phosphoprotein analyses revealed that even in these rare cases, oncogenic STAT5- or ERK-activation were mutually exclusive and segregated to competing clones. STAT5 and ERK engaged opposing biochemical and transcriptional programs orchestrated by MYC and BCL6, respectively. Genetic reactivation of the divergent (suppressed) pathway came at the expense of the principal oncogenic driver and reversed malignant transformation. Conversely, Cre-mediated deletion of divergent pathway components triggered leukemia-initiation and accelerated development of fatal disease. Thus, persistence of divergent signaling pathways represents a powerful barrier to malignant transformation while convergence on one principal driver defines a key event during leukemia-initiation. Proof-of-concept studies in patient-derived B-ALL cells revealed that pharmacological reactivation of suppressed divergent circuits strongly synergized with direct inhibition of the principal oncogenic driver. Hence, pharmacological reactivation of divergent pathways can be leveraged as a previously unrecognized strategy to deepen treatment responses and to overcome drug-resistance. Current treatment approaches for drug-resistant cancer are focused on drug-combinations to suppress the central oncogenic driver and multiple alternative pathways^14-17^. Here, we introduce a concept based on inhibition of the principal driver combined with pharmacological reactivation of divergent pathways.

---

During the earliest stages of B-cell development, cytokine receptors (e.g. IL7R, CRLF2) initiate survival signals by phosphorylation of JAK2 and STAT5^2,3^. After productive rearrangement of immunoglobulin V region genes and expression of a pre-B cell receptor (pre-BCR), an additional set of survival and proliferation signals involves the pre-B cell linker BLNK and phosphorylation of ERK kinases^8,18^. Mirroring their significance in normal B-cell development, STAT5- and ERK-mediated survival signals are frequently mimicked by transforming oncogenes in B-cell acute lymphoblastic leukemia (B-ALL). For instance, lesions in cytokine receptor genes (*IL7R, CRLF2, EPOR, PDGFRA, PDGFRB*)^4^ and cytokine receptor-associated JAK (*JAK1, JAK2, JAK3*)^5^ and ABL1 tyrosine kinases (*BCR-ABL1, ETV6-ABL1, NUP214-ABL1*)^6^ induce oncogenic STAT5-signaling and mimic constitutively active cytokine receptors. Likewise, activating lesions of RAS-pathway genes (*NRAS, KRAS, PTPN11, NF1*)^9^ and *BRAF*^10^ cause oncogenic ERK signaling, which acts as a functional mimic of pre-BCR-signaling^8,11,13^. Malignant transformation typically involves multiple genetic lesions whose combined activity gives rise to cancer. From this perspective, one would predict that addition of more oncogenic drivers to an existing set of mutations will invariably accelerate tumor progression. However, recent findings in B cell malignancies suggest that this is not always the case. For instance, loss of the tumor suppressor *PTEN* (resulting in hyperactivation of PI3K signaling) was synthetic lethal in BCR-ABL1 and NRAS^G12D^-driven B-ALL^19^. Likewise, loss-of-function of the tumor suppressors IRF4 and SPIB was incompatible with the oncogenic MYD88 driver in diffuse large B-cell lymphoma^20^. Here we studied the interaction of STAT5- and ERK-signaling pathways during normal B-cell development and malignant B-cell transformation.

## Activating mutations of STAT5- and ERK-pathways are frequent but mutually exclusive in B-ALL

Studying genetic lesions in 1,148 cases of B-ALL from pediatric and adult clinical cohorts^21-25^ (**Supplementary Tables 1-4**), we found STAT5-activating lesions (detailed in **Supplementary Table 1**) in 372 cases (31.4%) and ERK-activating lesions (detailed in **Supplementary Table 1**) in 386 cases (33.6%). Of the 721 cases (62.8%) that carried a driver mutation in at least one of the two pathways, only 37 (3.2%) had activating mutations in both pathways, suggesting that concurrent activation of STAT5- and ERK-occurs much less frequently in B-ALL than expected by chance (odds ratio 0.13; *P*=2.2e-16; **Fig. 1a**).

**Figure 1:**
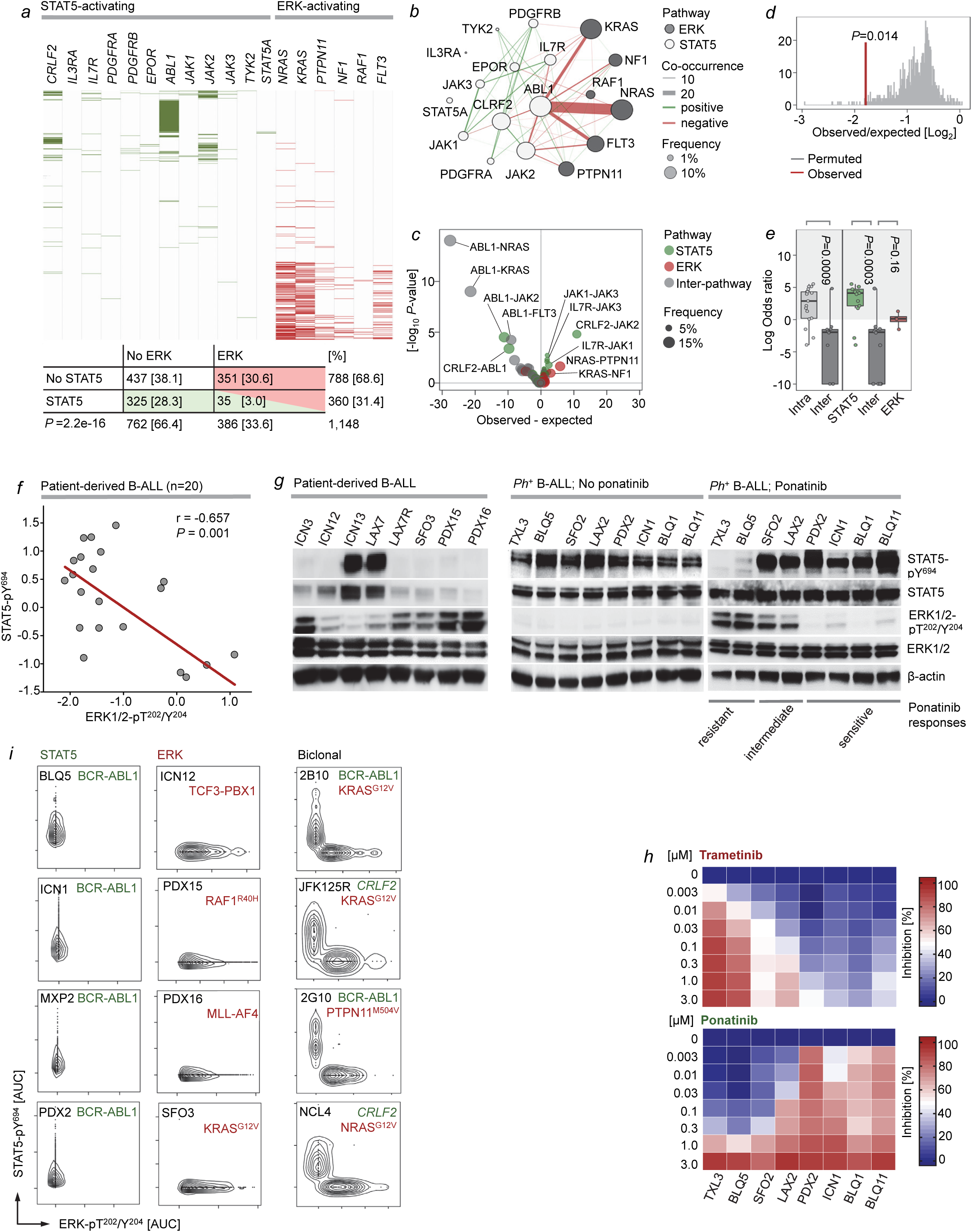
Segregation of STAT5- and ERK-activating oncogenic lesions in human B-ALL. **(a)** STAT5- and ERK-pathway mutations were studied in 1,148 patient-derived B-ALL samples (**Supplementary Table 1**). Null hypothesis is that STAT5- and ERK-pathway mutations occur independently of each other. The expected co-occurrence of the two mutations under the null hypothesis was 125. The observed co-occurrence of the two mutations was 37, significantly lower than the expected (*P*<2.2e^-16^, Fisher’s exact test). (**b**) To analyze gene-gene co-occurrence in a more unsupervised manner, Fisher’s test was run for each gene-pair and plotted as a co-occurrence network – pathway assignment for each gene is indicated by node color (grey = STAT5, white = ERK), and mutation frequency by node size. Direction of Fisher’s result is indicated by line color (green = positive/greater than expected, red = negative/less than expected), line width represents strength of association (–log10 p-value * |logOR|). OR: odds ratio. (**c**) Volcano plot of gene-gene co-occurrence results. Each point represents a gene-pair, colored by pathway assignment for the pair (green = both STAT5, red = both ERK, gray = interpathway), selected gene pairs are labeled. (**d**) As an alternative method to test for overall difference in co-occurrence between pathways, Fishers test was run over 10, 000 permutations with random shuffling of gene-pathway assignments on each iteration to generate null distribution for the hypothesis that pathway does not affect co-occurrence, and observed value was compared. (**e**) Distribution on logOR values from Fisher’s test grouped by gene-pair pathway assignment. Overall shifts in odds ratio are tested by Welsh t-test (left panel; *P*=0.0009), Tukey HSD (*P*=0.0003 and *P*=0.16, right panel) and ANOVA (overall; *P*=0.0004). Low frequency, non-significant gene-pairs were excluded to avoid extreme odds ratios from biasing results. (**f**) Western blot analyses were performed to evaluate correlation between phospho-ERK-T^202^/Y^204^ and phospho-STAT5-Y^694^ levels as determined by densitometry in patient-derived B-ALL cells (n=20). Shown is the correlation plot (*P*=0.001, two-tailed *t*-test; *r*=-0.657, Pearson r). (**g**) Levels of phospho-STAT5-Y^694^, STAT5, phospho-ERK-T^202^/Y^204^ and ERK were assessed by Western blot in patient-derived B-ALL cells (n=8, left), BCR-ABL1 B-ALL PDX before (n=8, middle) and after intermittent treatment with ponatinib (n=8, right). (**h**) Patient-derived *Ph*^+^ B-ALL cells were treated with increasing concentrations of trametinib or ponatinib. Percentage growth inhibition at each concentration of trametinib (top) and ponatinib (bottom) is presented (means of 3 independent experiments). (**i**) Single-cell phosphoprotein analyses for phospho-STAT5-Y^694^ and phospho-ERK-T^202^/Y^204^ were performed for patient-derived B-ALL samples. The scWest chips were first simultaneously probed with phospho-STAT5-Y^694^ and phospho-ERK-T^202^/Y^204^ antibodies, followed by fluorescent secondary antibodies. To confirm identified peaks were associated with cell occupancy, scWest chips were then probed for Histone H3, followed by TOTO-1 DNA staining.

Mutual exclusivity of mutations could also reflect their functional redundancy when they affect the same pathway or promote survival and proliferation in similar ways. In this case, underrepresentation of cooccurrence reflects lack of selective advantage rather than interference between the two lesions. Hence, if there was similar exclusivity of lesions within the STAT5 and ERK pathways (intra-pathway) as across pathways (inter-pathway), this likely suggests redundancy rather than interference between two lesions. We therefore performed an unsupervised analysis of mutational co-occurrence between all lesion pairs to determine whether inter-pathway STAT5-ERK mutation combinations within the same patient were more strongly selected against than exclusivity of redundant of intra-pathway driver mutations. Indeed, we observed overall significantly stronger exclusivity between inter-pathway lesions compared to intra-pathway lesions, with the strongest underrepresentation of co-occurrence of *ABL1* with *NRAS* and *KRAS* lesions (**Fig. 1b-c**). Permutation of overall pathway-exclusivity with random re-assignment of lesions to pathways indicates that concurrent mutation of multiple drivers is indeed counter-selected regardless of pathway, but that the observed STAT5-ERK pathway co-occurrence is significantly less frequent than this empirical distribution (*P*=0.014; **Fig. 1d**); this is further supported by significantly lower log odds ratios for inter- vs intra-pathway driver combinations (*P*=0.001; **Fig. 1e**).

Activating STAT5- and ERK-pathway lesions are also common in acute myeloid leukemia (AML). To determine whether segregation of STAT5- and ERK-pathway lesions represents a unique feature of B-ALL, we studied co-occurrence of STAT5- and ERK-pathway lesions in 916 patient-derived AML samples. STAT5- (100 cases; 10.9%) and ERK- (309 cases; 33.7%) pathway lesions were frequent in AML, however, co-occurrence was only found in 24 cases (2.6%; **Supplementary Table 2**). As in B-ALL, the observed co-occurrence of STAT5- and ERK-pathway lesions in AML was significantly below random co-occurrence (odds ratio 0.59, *P=*0.033, Fisher’s exact test; **Extended Data Figure 1a**). Unlike in B-ALL, however, mutual exclusivity of inter-pathway pairs was not significantly stronger than the observed mutual exclusivity between multiple intra-pathway drivers (*P*=0.63; **Extended Data Figure 1b-e**). Hence, while STAT5- and ERK-activating mutations are frequent in both B-ALL and AML, driver mutations in the two pathways are strongly segregated in the former but not in the latter.

## Oncogenic activation of STAT5- and ERK-signaling is mutually exclusive in B-ALL

In addition to segregation of STAT5- and ERK-activating mutations, also biochemical activation of STAT5- and ERK-signaling as well as STAT5- and ERK-dependent drug-responses followed a divergent and mutually exclusive pattern: Western blot analyses of 20 B-ALL patient-derived xenografts (PDX) established in our laboratory (**Supplementary Table 5**) confirmed an inverse relationship between STAT5-pY^694^ and ERK-T^202^/Y^204^ phosphorylation (**Fig. 1f**). The tyrosine kinase inhibitor (TKI) ponatinib suppresses STAT5-signaling in B-ALL driven by the BCR-ABL1 kinase. Intermittent treatment of 8 BCR-ABL1-driven B-ALL with ponatinib over three weeks resulted in gradual development of TKI-resistance in 4 cases, while the other 4 cases remained sensitive. Two of the ponatinib-resistant cases (TXL3 and BLQ5) lost STAT5-activity and instead acquired *de novo* phosphorylation of ERK (**Fig. 1g**). Consistent with ponatinib-resistance and a switch from STAT5-phosphorylation to *de novo* ERK-phosphorylation, TXL3 and BLQ5 cells also acquired sensitivity to trametinib, a small molecule inhibitor of MAP2K1 (MEK), an essential upstream activator of ERK (**Fig. 1h**).

## Single-cell mutation and phosphoprotein analyses reveal clonal segregation of STAT5 and ERK-activation

To test whether STAT5- and ERK-activating mutations in rare cases with dual pathway activation co-occurred in the same cells or were segregated to distinct clones, we studied these mutations at the single-cell level based on two approaches, namely single-cell amplicon sequencing (**Supplementary Table 4**) and single-cell phosphoprotein analyses for STAT5-Y^694^ and ERK-T^202^/Y^204^ (**Fig. 1i**). Immunodeficient NSG mice (n=30) bearing PDX from 9 patients of *Ph*^+^ B-ALL (STAT5-driven) were treated with ponatinib. After initial remission, mice relapsed with fatal B-ALL in 24 cases. Reflecting strong selective pressure, genomic analyses revealed that the relapsed B-ALLs acquired additional STAT5- or ERK-activating lesions that ultimately subverted ponatinib-activity (**Supplementary Table 3**). In 8 cases, additional STAT5-activating mutations were acquired. In 13 cases, ponatinib-resistant clones emerged with ERK-activating lesions and in 3 cases, both STAT5- and ERK-activating lesions were found (**Supplementary Tables 3-4**). We selected these three PDX (1F10, 2B10, 2G10) for single-cell amplicon sequencing. To this end, single cells were sorted by flow cytometry from each PDX and subjected to rounds of nested PCR amplification and subsequent sequencing of cloned PCR products (**Supplementary Table 4**). In all three cases, target loci (*ABL1, KRAS, PTPN11*) were successfully amplified and sequenced from 71 cells (1F10), 79 cells (2B10) and 63 cells (2G10). STAT5- (*ABL1*) and ERK- (*PTPN11, KRAS*) activating mutations were amplified individually from single sorted B-ALL cells. However, mutant *ABL1* alleles were only co-amplified with wildtype *KRAS* and *PTPN11* alleles and *vice versa* (**Supplementary Table 4**). These results demonstrate that even in the few cases that harbored activating mutations in both STAT5- and ERK-pathways, these mutations were segregated to distinct clones. To corroborate pathway segregation at the level of STAT5- and ERK-phosphorylation, we performed single-cell phosphoprotein analyses for STAT5-pY^694^ and ERK-pT^202^/Y^204^ for patient-derived B-ALL samples with evidence of dual activation of STAT5 and ERK pathways (**Fig. 1i; Extended Data Figure 2**). Consistent with a scenario in which not only genetic STAT5- and ERK-activating lesions but also biochemical pathway activation was segregated to distinct clones, the single-cell phosphoprotein analysis revealed that STAT5- and ERK-phosphorylation was mutually exclusive and confined to separated, potentially competing clones (**Fig. 1i; Extended Data Figure 2**). The few events (∼2%) in which concurrent phosphorylation of STAT5 and ERK was detected may be attributed to a similar low frequency of two cells being loaded and analyzed in the same well (**Extended Data Figure 2**).

## Reciprocal suppression of oncogenic STAT5- and ERK-signaling results in pathway convergence

Unsupervised mutation analyses in AML revealed that mutual exclusivity between STAT5- and ERK-pathways was not different between mutations within the same pathway, reflecting functional redundancy rather than negative interference. In B-ALL, however, we observed much stronger inter-than intra-pathway mutual exclusivity, indicating functional incompatibility and negative selection of concurrent STAT5- and ERK-activation. To determine the mechanistic basis of segregation of STAT5- and ERK-activation, we examined whether inducible activation of one pathway affected biochemical activity of the other. We developed inducible systems for STAT5- and ERK-activation and measured the impact on phosphorylation of Stat1, Stat3 and Stat5 proteins as well as MAP kinases Erk, p38α and Jnk (**Figure 2a**). Inducible expression of a constitutively active mutant of Stat5a in IL7-dependent pro-B cells strongly increased Stat5-phosphorylation (Y^694^) and almost entirely abolished Erk-phosphorylation (T^202^/Y^204^; **Fig. 2a**). Like ERK, also JNK- and p38α-activity was suppressed by STAT5. However, only ERK but not JNK and p38α, was phosphorylated upon inducible activation of oncogenic NRAS^G12D^ in IL7-dependent pro-B cells. STAT1-phosphorylation was barely detectable in B-ALL cells. Interestingly, STAT3 was strongly phosphorylated downstream of RAS-activation on both Y^704^ and S^727^. Phosphorylation of these sites was suppressed by oncogenic STAT5 signaling (**Fig. 2a, Extended Data Figure 3a**). In contrast to increased phosphorylation of Stat3, inducible expression of oncogenic NRAS^G12D^ resulted in dephosphorylation of Stat5 (**Fig. 2a**), suggesting that STAT3 and STAT5 oppose each other.

**Figure 2:**
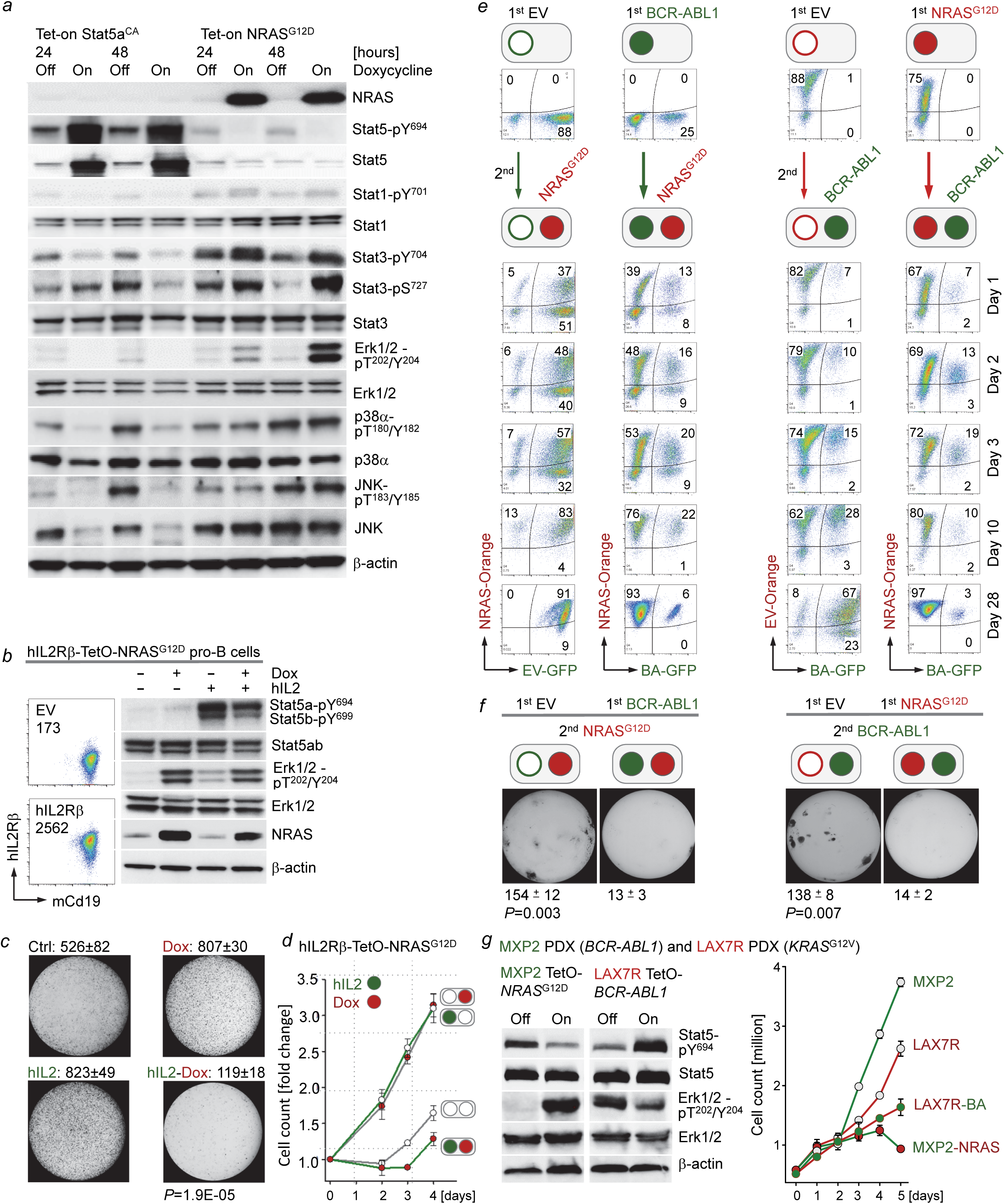
Concurrent oncogenic STAT5- and ERK-activation subverts B-cell leukemogenesis. **(a)** Levels of NRAS, Stat5-pY^694^, Stat5, phospho-Stat1-Y^701^, Stat1, phospho-Stat3-Y^704^, phospho-Stat3-S^727^, Stat3, Erk-pT^202^/Y^204^, Erk1/2, p38α-pT^180^/Y^182^, p38α, JNK-pT^183^/Y^185^ and JNK were assessed by Western blotting following Dox-induced expression of Stat5a^CA^ or *NRAS*^G12D^ in IL7-dependent mouse pro-B cells. (**b**) IL7-dependent Tet-on NRAS^G12D^ mouse pro-B cells expressing hIL2Rβ were induced with Dox (24 hr), hIL2 (15 minutes, 50 ng/µL) or a combination of both. Levels of Stat5-pY^694^, Stat5, Erk-pT^202^/Y^204^ and Erk were assessed by Western blotting. FACS analysis was performed to measure surface expression of hIL2Rβ and mCd19. (**c-d**) IL7-dependent hIL2Rβ-Tet-on NRAS^G12D^ mouse pro-B cells were induced with Dox, hIL2 (10 ng/µL), or a combination of both. 50,000 cells were seeded in methylcellulose for colony formation assays (10 days, n=3, **c**). Viable cell counts (**d**) were measured at various time points. Dox and hIL2 were replenished every other day. Shown are average values from 3 independent experiments. (**e**) IL7-dependent B-cell precursors were retrovirally transduced with EV, *BCR-ABL1*-GFP (BA-GFP), EV-Orange, or NRAS^G12D^-Orange. Following the first round of transductions, cells were transduced with NRAS^G12D^-Orange or *BCR-ABL1*-GFP as indicated. Flow cytometry was performed to monitor the proportions of GFP^+^, Orange^+^, and double-positive cells at various time points following transductions. Representative FACS plots from 3 independent experiments. (**f**) Cells from (**e**) were sorted for double-positive (GFP^+^ and Orange^+^) populations, and 10, 000 cells were seeded in methylcellulose for colony formation assays (10 days, n=3). *P*=0.003 (left panel) and *P*=0.007 (right panel; two-tailed *t*-test). (**g**) Patient-derived BCR-ABL1 B-ALL cells (MXP2) expressing Tet-on NRAS^G12D^ and patient-derived *KRAS*^G12V^ B-ALL cells (LAX7R) expressing Tet-On *BCR-ABL1* were induced with Dox. Levels of STAT5-pY^694^, STAT5, ERK-pT^202^/Y^204^ and ERK (left) as well as viable cell counts (right) were measured upon induction with doxycycline (Dox).

To confirm biochemical interference between STAT5- and ERK-signaling in an independent experimental model, we engineered murine pro-B cells to express (i) the human IL2 receptor β (hIL2Rβ) chain which enables phosphorylation of Stat5a-Y^694^ and Stat5b-Y^699^ in response to human IL2-binding and (ii) doxycycline-inducible NRAS^G12D^ (**Fig. 2b**). Tightly controlled activation of Stat5 (by addition of human IL2), Erk (by induction of NRAS^G12D^) or a combination of both, confirmed biochemical interference between activation of STAT5 and ERK pathways (**Fig. 2b**). Plating of hIL2Rβ TetO-NRAS^G12D^ pro-B cells revealed increased colony formation by activation of Stat5 (hIL2) or Erk (Dox) alone, which was reduced by ∼7-fold upon dual Stat5-Erk pathway activation (**Fig. 2c**). Likewise, activation of Stat5 or Erk alone stimulated proliferation of hIL2Rβ TetO-NRAS^G12D^ pro-B cells, while concurrent activation had a strong growth-suppressive effect (**Fig. 2d**). These observations indicate that signal input from divergent signaling pathways represents a powerful barrier against malignant transformation.

The analysis of cooccurrence networks of STAT5- and ERK-activating lesions identified a particularly strong negative association between *BCR-ABL1* and oncogenic *NRAS* lesions (**Fig. 1b-c**). For this reason, we modelled concurrent activation of BCR-ABL1 with NRAS^G12D^ (**Fig. 2e-f**). IL7-dependent pro-B cells were initially transduced with either BCR-ABL1-GFP (BA-GFP) or NRAS^G12D^-Orange (NRAS-Orange) or empty vector controls (EV-GFP or EV-Orange). One week later, BA-GFP B-ALL cells were transduced with NRAS-Orange for concurrent activation of Erk and NRAS-Orange B-ALL cells were transduced with BA-GFP for concurrent activation of Stat5. The ability of oncogenic Stat5 (GFP^+^) and Erk (Orange^+^) signaling to contribute to the dominant clone was monitored by flow cytometry over time. When introduced as sole oncogenic driver following transduction with neutral EV-Orange or EV-GFP, both Stat5 (EV-Orange +BA-GFP) and Erk (EV-GFP +NRAS-Orange) gave rise to large double-positive populations. In contrast, when introduced as secondary oncogene engaging a diverging pathway from the principal driver, the frequency of double-positive cells was dramatically reduced for both Stat5 (from 67% following EV to 3% following Erk) and Erk (from 91% following EV to 6% following Stat5; **Fig. 2e**). The ∼15-fold reduction of double-positive cells for secondary divergent oncogenes was mirrored by a similar (∼10-fold) reduction of colony formation comparing single- and dual pathway activation (**Fig. 2f**). Hence, concurrent activation of Stat5 and Erk was not productive and reduced competitive fitness of double-positive clones, unless the conflict was resolved and clones converged on one pathway (single-positive cells). In the experiments that were performed, the conflict between Stat5 and Erk pathways was eventually resolved in favor of a dominant Erk-driven clone. Even when Stat5 was introduced first, ultimately Erk (Orange^+^) clones prevailed. Clonal dominance of Erk-over Stat5-driven B-ALL in these experiments mirrored findings from recent work that traced variant allele frequencies of STAT5- and ERK-activating mutations in 17 B-ALL patients^26^. The competitive advantage of ERK-over STAT5-activation was confirmed in two clonal outgrowth experiments based on (i) murine pro-B cells transduced with either STAT5-GFP or KRAS^G12V^-mCherry and (ii) competition between *IL7R*-mutant (STAT5) and *KRAS*^G12V^ (ERK)-driven B-ALL PDX cells from the same patient (**Extended Data Figure 3b-c**).

To confirm pathway interference between BCR-ABL1 and oncogenic RAS (**Fig. 1b-c**) in patient-derived B-ALL cells, we engineered PDX from *BCR-ABL1*- (MXP2) and *KRAS*^G12V^- (LAX7R) B-ALL with doxycycline-inducible constructs for expression of NRAS^G12D^ and BCR-ABL1, respectively (**Fig 2g-h, Extended Data Figure 3d-i**). Concurrent activation of a divergent pathway suppressed biochemical activity of the principal oncogenic driver and proliferation (**Fig. 2g**), which was paralleled by cellular senescence and cell death. Dual pathway activation resulted in a senescent phenotype (senescence-associated β-galactosidase activity, upregulation of p16 and p21) and ultimately apoptosis and cell death (Annexin V^+^; **Extended Data Figure 3d-i**).

## Genetic deletion of divergent pathway components enables malignant transformation and leukemia-initiation

Compared to activation of oncogenic STAT5 or ERK alone, concurrent activation of both pathways caused reciprocal interference of signaling, compromised colony formation and ultimately induced cellular senescence and apoptosis (**Fig. 2, Extended Data Figure 3d-i**). Hence, we tested the hypothesis that background signaling from pathways that diverge from the principal oncogenic driver represents a barrier against malignant transformation. To this end, we developed mouse models for STAT5- and ERK-driven B-ALL, in which tamoxifen-inducible activation of Cre resulted in genetic ablation of *Stat5* (*Stat5*^fl/fl^) or Erk2 (*Mapk1*^fl/fl^). Consistent with reciprocal biochemical cross-inhibition, inducible deletion of *Mapk1* (Erk2) resulted in loss of Stat3-S^727^ but increased Stat5-phosphorylation. Conversely, inducible *Stat5*-deletion increased phosphorylation of both Erk and Stat3-S^727^ (**Fig. 3a-b**). As expected, genetic ablation of the principal oncogenic pathway (*Stat5*^fl/fl^ in BCR-ABL1- and *Mapk1*^fl/fl^ in NRAS^G12D^-driven B-ALL) induced rapid depletion of cells from cell culture in growth competition assays (**Fig. 3c-d**). In contrast, Cre-mediated deletion of secondary divergent pathways, namely *Mapk1* in Stat5-driven and *Stat5* in Erk-driven B-ALL, transiently slowed cell growth but did not result in substantial depletion of cells from cell culture (**Fig. 3c-d**). Compared to B-ALL cells that retained intact divergent pathway components, Cre-mediated deletion strikingly increased colony formation in secondary and tertiary replatings (**Fig. 3e-f**). To examine potential effects of deletion of divergent pathway components on B-ALL leukemia-initiation, we performed limiting dilution transplant experiments based on 100, 1,000 and 10,000 pro-B cells that carried Stat5- or Erk-driver oncogenes and *Mapk1*^fl/fl^ (Erk) and *Stat5*^fl/fl^ alleles for inducible ablation of divergent pathways. *Mapk1*^fl/fl^ pro-B cells with BCR-ABL1 (Stat5) and *Stat5*^fl/fl^ pro-B cells with NRAS^G12D^ (Erk) were studied either carrying EV, for retention, or tamoxifen-inducible Cre, for deletion of divergent pathway components. Five days after injection and engraftment, NSG recipient mice were injected with tamoxifen for activation of Cre or EV controls. At all three dose levels, deletion of *Mapk1* accelerated the onset of overt Stat5-driven B-ALL. Likewise, deletion of *Stat5* accelerated leukemia-initiation in Erk-driven B-ALL. At the two lower dose levels (1,000 and 100 cells), deletion of *Stat5* was even required for initiation of Erk-driven B-ALL. Injection of 100 or 1,000 Erk-driven B-ALL cells failed to initiate fatal disease in transplant recipients unless *Stat5* was deleted. Inducible deletion of *Stat5* prompted the onset of lethal Erk-driven B-ALL at 100% penetrance (**Fig. 3g-h**). These results were further corroborated in experiments based on pharmacological inhibition of divergent pathways: Colony formation of murine pro-B cells carrying an inducible *BCR-ABL1* knockin allele was substantially increased by Trametinib-mediated ablation of ERK-signaling. Likewise, colony formation of murine pro-B cells carrying an inducible *KRAS*^G12D^ knockin allele (**Supplementary Table 6**) was increased by ∼6-fold when divergent Stat5 signaling was suppressed by Ruxolinitib (**Extended Data Figure 4a-b**). Together, these results indicate that ablation of divergent signaling and convergence on one principal oncogenic driver represents a critical event during early stages of B cell transformation and leukemia-initiation. In agreement with deletion of *Mapk1* as a leukemia-initiating event in Stat5-driven B-ALL, a recent Sleeping Beauty transposon screen suggested that truncation of the ERK-signaling activator *Sos1* can precipitate the onset of STAT5-driven B-ALL^27^. Importantly, recent observations in patients suggest that prolonged pharmacological suppression of JAK2-STAT5 signaling by Ruxolitinib could prime dormant B-cell clones to develop overt ERK-driven B-cell lymphoma^28^.

**Figure 3:**
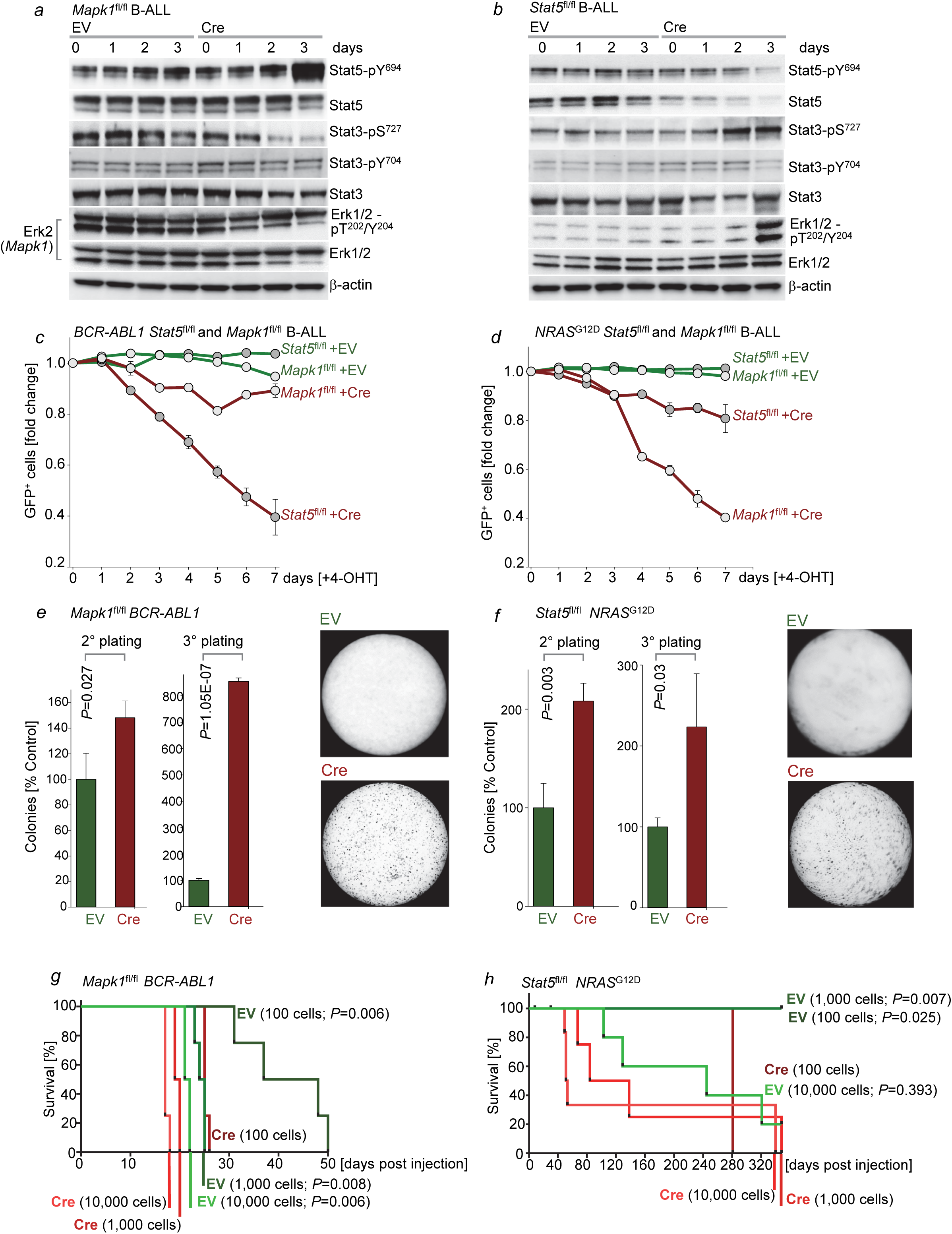
Genetic deletion of alternative pathways triggers STAT5- and ERK-driven leukemia-initiation. **(a)** Levels of phospho-Stat5-Y^694^, Stat5, phospho-Stat3-S^727^, Stat3, phospho-Erk1/2-T^202^/Y^204^ and Erk1/2 were measured by Western blotting in *BCR-ABL1*-driven B-ALL cells upon Cre-mediated ablation of *Stat5*. **(b)** Levels of phospho-Stat5-Y^694^, Stat5, phospho-Stat3-S^727^, Stat3, phospho-Erk1/2-T^202^/Y^204^ and Erk1/2 in *NRAS*^*G12D*^-driven B-ALL cells upon Cre-mediated deletion of *Mapk1 (Erk2)*. (**c**) *Stat5*^fl/fl^ or *Mapk1*^fl/fl^ *BCR-ABL1* B-ALL cells were transduced with ER^T2^-GFP or Cre-ER^T2^-GFP. Enrichment or depletion of GFP^+^ cells was monitored by flow cytometry upon 4-OHT induction (n=3). (**d**) *Stat5*^fl/fl^ or *Mapk1*^fl/fl^ *NRAS*^G12D^ B-ALL cells were transduced with ER^T2^-GFP or Cre-ER^T2^-GFP. Flow cytometry was performed to measure enrichment or depletion of GFP^+^ cells upon 4-OHT induction (n=3). (**e and f**) *Mapk1*^fl/fl^ *BCR-ABL1*-driven B-ALL cells (**e**) and *Stat5*^fl/fl^ *NRAS*^G12D^-transformed B-ALL cells (**f**) were sorted for GFP^+^ populations, and 10, 000 sorted cells were seeded in semi-solid methylcellulose in the presence of 4-OHT and monitored for colony formation for 10-14 days. Serial re-plating was performed (n=3). *P*-values shown were determined by two-tailed *t*-test. (**g and h**) Kaplan-Meier analyses (Mantel-Cox log-rank test) of recipient mice (n=4 per group) injected with *BCR-ABL1*-(**g**) or *NRAS*^G12D^-driven (**h**) B-ALL cells following 4-OHT-induced deletion of *Mapk1* (**g**; 24 hr) or *Stat5* (**h**; 24 hr).

## STAT5- and ERK-activation define distinct stages of early B-cell development

Activation of STAT5 (downstream of cytokine receptors in pro-B cells) and ERK (downstream of the pre-BCR in pre-B cells) are linked to distinct stages of early B-cell development that are separated by rearrangement of immunoglobulin heavy (Ig-HC) and light chain (Ig-LC) genes^3,18,29-30^. To test whether STAT5- and ERK-activating lesions in patient-derived B-ALL cells are linked to cellular origins from distinct stages of B-cell development, we studied the configuration of Ig-κ (*IGKV*; 2p11) and λ- (*IGLV*; 22q11.2) light chain loci and Ig-LC surface expression by flow cytometry in 22 cases of B-ALL with STAT5- and 42 cases with ERK-activating lesions (**Extended Data Figure 5a-b**). Even though aberrant RAG1/RAG2-mediated recombinase activity in B-ALL cells^31^ can randomly target Ig-LC loci, the analysis revealed a significant association between STAT5-driven B-ALL and germline configuration on one and ERK-driven B-ALL and rearranged Ig-LC loci on the other hand (*P*=0.008; **Extended Data Figure 5a-b**). Inducible activation of a gain-of-function variant of Stat5a (TetO-Stat5a^CA^) in *BCR-ABL1* B-ALL cells induced upregulation of cytokine receptor-related genes, while inducible ablation of Stat5 (*Stat5*^fl/fl^) in *BCR-ABL1* B-ALL cells resulted in loss of expression of these genes (**Extended Data Figure 5c-d**). Conversely, genetic ablation of *Stat5* upregulated the pre-B cell linker Blnk and Ptpn6, while these genes were transcriptionally repressed by Stat5a^CA^. Negative regulation of Blnk by Stat5 was confirmed by Western blot at the protein level based on gain and loss-of-function experiments (**Extended Data Figure 5e**). Studying gene expression in patient-derived B-ALL samples from the P9906 clinical trial, B-ALL samples with STAT5-activating lesions expressed immature pro-B cell antigens (CD43, KIT) and cytokine receptor components (IL7R, CRLF2) at high levels. In contrast, B-ALL samples carrying ERK-activating mutations expressed higher levels of Ig-LC genes (IGLV3-19, IGLV-1-44, IGLC2), the pre-B cell linker Blnk and PTPN6, a negative regulator of STAT5 **(Extended Data Figure 5f-h**).

Analyzing 17 B-ALL PDX as well as genetic B-ALL mouse models studied here by flow cytometry, we further corroborated the association between oncogenic STAT5-signaling and a pro-B cell phenotype (CD34^+^) and higher levels of cytokine receptor expression (CRLF2, IL7R), while ERK-driver lesions were associated with a pre-B cell phenotype (VPREB1, CD21, IgM, Ig-LC; **Extended Data Figure 5b** and **5i-n**). Likewise, in mouse models for STAT5- and ERK-driven B-ALL, we observed that STAT5-activation was associated with a pro-B cell phenotype (Cd43^+^, Il7r^high^, Cd21^-^, pre-BCR^-^, Ig-LC^-^), whereas ERK-activation was linked to a more mature pre-B cell phenotype (Cd43^-^, Il7r^Low^, Cd21^+^, pre-BCR^+^, Ig-LC^+^**; Extended Data Figure 5o**).

## MYC and BCL6 as downstream effectors of STAT5- and ERK-pathways

Hence, we examined how the pro-B to pre-B cell transition and Ig-gene rearrangement at the pre-BCR checkpoint affects permissiveness to oncogenic STAT5- and ERK-signaling. Since STAT5- and ERK-lesions were associated with pro-B cell and pre-B cell phenotypes, respectively, we studied normal activation of the two pathways at the pro-B to pre-B cell transition during early B-cell development. Studying Stat5- and Erk-activation by single cell phosphoprotein analysis in sorted bone marrow pro-B cell (Hardy Fractions B and C) and pre-B cell/immature B-cell (Hardy Fractions E-F) populations, activation of Stat5 during pro-B cell stages was turned off at the pre-B cell receptor (pre-BCR) checkpoint and switched to Erk activation in more mature B-cell development (**Figure 4a**).

**Figure 4:**
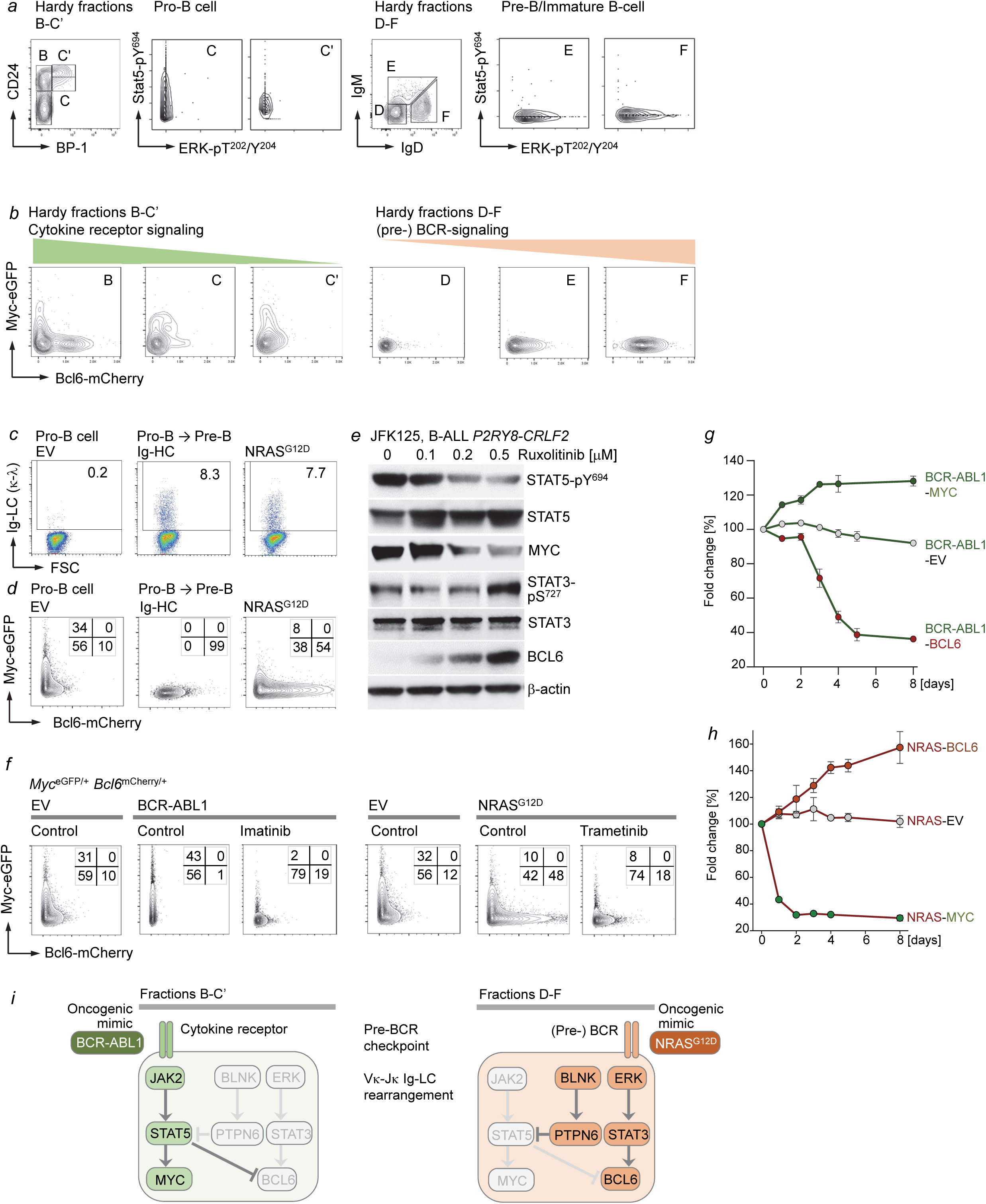
STAT5-MYC and ERK-BCL6 signaling are incompatible and define distinct stages of B cell development. **(a)** Hardy Fractions B-F analysis was performed with *Myc*^eGFP/+^ *Bcl6*^mCherry/+^ bone marrow cells (n=3). Single-cell Western blot analyses for phospho-Stat5-Y^694^ and phospho-Erk-T^202^/Y^204^ were performed for the indicated fractions. (**b**) Hardy Fraction B-F analysis was performed with *Myc*^eGFP/+^ *Bcl6*^mCherry/+^ bone marrow cells. Expression of eGFP and mCherry was monitored by flow cytometry in each fraction (B-F; n=3). (**c**) Surface expression of Igκ light chain (LC) was measured by flow cytometry in IL7-dependent pro-B cells (left), and upon Dox-inducible expression of μHC (middle) or NRAS^G12D^ (right). Shown are representative results from 3 independent experiments. (**d**) IL7-dependent *Myc*^eGFP/+^ *Bcl6*^mCherry/+^ B-cell precursors were induced to differentiate or were transduced with NRAS^G12D^. Expression of eGFP and mCherry was measured by flow cytometry. Shown are representative results from 3 independent experiments. (**e**) Patient-derived *Ph*-like B-ALL cells were treated with increasing concentrations of ruxolitinib (µM), and Western blot analyses were performed to assess levels of phospho-STAT5-Y^694^, STAT5, MYC, phospho-STAT3-S^727^, STAT3 and BCL6. (**f**) *Myc*^eGFP/+^ *Bcl6*^mCherry/+^ B-cell precursors were transduced with EV, *BCR-ABL1* or *NRAS*^G12D^, and were subsequently treated with vehicle control, imatinib (1 µM) or trametinib (10 nM) as indicated. Expression of eGFP and mCherry was monitored by flow cytometry. Shown are representative results from 3 independent experiments. (**g**) *BCR-ABL1*-driven B-ALL cells were transduced with EV control, GFP-tagged MYC or GFP-tagged BCL6. Enrichment or depletion of GFP^+^ populations were monitored by flow cytometry (n=3). (**h**) *NRAS*^G12D^-transformed B-ALL cells were transduced with EV control, GFP-tagged MYC, or GFP-tagged BCL6. Enrichment or depletion of GFP^+^ populations were monitored by flow cytometry (n=3). (**i**) Like in normal B cell development, B-ALL subtypes can be traced to specific differentiation stages with distinct requirements for survival and proliferation signals. For instance, pro-B cells depend on cytokine receptor signaling and activation of STAT5 but not ERK. Conversely, pre-B cells depend on pre-BCR signaling and activation of ERK but not STAT5. STAT5-driven B-ALL cells depend on oncogenic mimics of cytokine receptor signaling and resemble pro-B cells that depend on STAT5-activation downstream of cytokine receptors. Mimicking pre-BCR signaling at the pro-B to pre-B cell-transition, oncogenic RAS-signaling suppresses STAT5-signaling and induces de novo expression of *BCL6*.

While Stat5-signaling transcriptionally activated Myc and suppresses Bcl6 (**Extended Data Figure 5c**), Erk-activity at the pre-BCR checkpoint induced downregulation of Myc with concurrent upregulation of Bcl6 (**Extended Data Figure 5p-q**). To identify Myc- and Bcl6-expressing populations during early B-cell development, we generated a B-cell-specific *Myc*-eGFP x *Bcl6*-mCherry double reporter knockin mouse model. Protein expression of Myc and Bcl6 in these populations was validated by cell sorting and Western blot analysis (**Extended Data Figure 5r-s**). Studying Myc^+^ and Bcl6^+^ cells during B-cell differentiation *in vivo* showed a similar transition from Myc^+^ Bcl6^-^ to Myc^-^ Bcl6^+^ stages. Cytokine-dependent pro-B and early pre-B cell subsets (Hardy fractions B-C’) included double-negative and Myc-expressing cells while BCR-dependent B-cells (Hardy fractions E-F) expressed almost exclusively Bcl6 (**Fig. 4b**). Late pre-B cells (Fraction D) that have already downregulated cytokine receptors but still undergo Ig-LC gene rearrangement to express a BCR mark the intersection between Myc^+^ Bcl6^-^ to Myc^-^ Bcl6^+^ stages and express neither Myc nor Bcl6 (**Fig. 4b**). We induced the pro-B to pre-B-cell transition experimentally by forced expression of a functional Ig-HC in pro-B cells, which then gave rise to Ig-LC gene rearrangement and surface expression (**Fig. 4c**). Interestingly, activation of NRAS^G12D^ and ERK recapitulated the pro-B to pre-B cell transition and had a very similar effect on Ig-LC surface expression (**Fig. 4c**). As observed for sorted pro-B and pre-B cell populations (**Fig. 4b**), experimental induction of pro-B to pre-B-cell transition, either by forced Ig-HC expression or activation of oncogenic Erk signaling (*NRAS*^G12D^), resulted in a phenotypic switch from Myc^+^ Bcl6^-^ pro-B cell to Myc^-^ Bcl6^+^ pre-B cell stages (**Fig. 4d**).

## STAT5-MYC and STAT3-BCL6 engage mutually exclusive transcriptional programs

In a STAT5-driven B-ALL PDX, pharmacological inhibition of STAT5 had similar effects and resulted in strong upregulation BCL6 at the expense of MYC. Consistent with opposing roles of Stat5 and Stat3 in B-cells^32^, downregulation of STAT5-was paralleled by increased STAT3-phosphorylation (**Figure 4e**). While oncogenic Stat5-signaling (BCR-ABL1) promoted strong activation of Myc and suppression of Bcl6, engagement of Erk-signaling (NRAS^G12D^) had the opposite effect and caused strong activation of Bcl6-mCherry while Myc-eGFP was reduced (**Fig. 4f**). Inhibition of oncogenic ERK-activation using a small molecule inhibitor of MAP2K1 (Trametinib) largely reduced activity of the Bcl6-reporter. Consistent with other observations suggesting that STAT5- and ERK-activating lesions are incompatible (**Fig. 1, Fig. 2**), flow cytometry analysis of Myc^eGFP^-Bcl6^mCherry^ dual reporter cells showed an L-shaped pattern (**Fig. 4f**) comparable to STAT5-ERK single-cell phosphoprotein analyses (**Fig. 1i**). Reflecting distinct stages of early B-cell differentiation, these findings suggest that Stat5-Myc^+^ and Erk-Bcl6^+^ cells represent non-overlapping populations and likely competing clones in B-ALL. Since oncogenic activation of kinases upstream of STAT5 (e.g. JAK2, BCR-ABL1) and STAT3 (e.g. ERK) are not compatible, we tested whether activation of the STAT5- and STAT3-downstream targets, MYC and BCL6, was also mutually exclusive. Consistent with competitive antagonism between STAT5 and STAT3^32^, In BCR-ABL1-driven B-ALL, we observed predominant activation of the STAT5-MYC pathway, while NRAS^G12D^ (ERK)-driven B-ALL mainly activates STAT3-BCL6 (**Fig. 2a, Extended Data Figure 3c**). To test whether the two pathways are compatible at the transcriptional level, we transduced BCR-ABL1-driven B-ALL (STAT5-MYC) with BCL6 and ERK-driven B-ALL (STAT3-BCL6) with MYC. Transduction of MYC introduced a survival advantage in BCR-ABL1-driven B-ALL (STAT5), whereas BCL6 caused depletion of cells from cell culture in growth competition assays (**Fig. 4g**). Likewise, transduction of BCL6 promoted growth advantage of NRAS^G12D^-driven B-ALL (STAT3), but NRAS^G12D^-driven B-ALL cells were not permissive to MYC and rapidly depleted from cell culture (**Fig. 4h**). These results delineate two mutually exclusive pathways and demarcate distinct stages of B-cell development: while cytokine receptor signaling in pro-B cells or its oncogenic mimics (e.g. BCR-ABL1) activate JAK2 and STAT5 to promote a MYC-driven transcriptional program, pre-BCR signaling in pre-B cells or its oncogenic mimics (e.g. NRAS^G12D^) activate ERK and STAT3 to engage BCL6-dependent transcription (**Fig. 4i**).

## Developmental rewiring from STAT5-to ERK-signaling at the pre-B cell receptor checkpoint

Activity of the pre-BCR checkpoint and rearrangement of Ig-LC genes represent main differences between pro-B and pre-B cell stages. Interestingly, genes that are implicated in pre-BCR checkpoint function and Ig-LC assembly are expressed at higher levels in ERK-compared to STAT5-driven B-ALL, consistent with a more mature phenotype in the former compared to the latter (**Extended Data Figure 5a-b, 5f-o)**. To determine whether expression levels of these genes (IGLV, IGHM, BLNK, BCL6) are predictive of poor or favorable clinical outcomes, we compared overall and relapse-free survival of patients with B-ALL (COG P9906) that were segregated into two groups based on higher vs lower than median mRNA levels (top vs. bottom half) for these genes. When analyzed across the entire cohort, clinical outcomes were not significantly different between patients in the top vs. bottom half based on mRNA levels of these genes. However, when the analysis was performed separately for STAT5-driven (n=26) and ERK-driven (n=67) B-ALL patients, higher than median mRNA levels of *IGLV, IGHM* and BLNK predicted favorable outcomes in patients with STAT5 B-ALL but poor clinical outcomes in patients with ERK B-ALL. BCL6 was found as a predictor of poor outcome in ERK B-ALL while no significant difference was observed for patients with STAT5 B-ALL (**Extended Data Figure 6a**). These findings suggest that phenotypic differences between STAT5- and ERK-driven B-ALL likely extend to divergent course of disease and clinical outcome. To experimentally test if developmental rewiring at the pre-BCR checkpoint affects permissiveness to oncogenic STAT5 or ERK signaling, we tested activation of these pathways in flow-sorted mouse bone marrow pro-B cells (CD19^+^ CD43^+^ Ig-HC^-^ VpreB^-^) and pre-B cells (CD19^+^ CD43^-^ Ig-HC^+^ VpreB^+^). While pro-B cells were permissive to BCR-ABL1 (STAT5), oncogenic activation of ERK by the pre-BCR mimic LMP2A11 induced cellular senescence and rapid cell death (**Extended Data Figure 6b-i**). Conversely, pre-B cells that have passed the pre-BCR checkpoint were permissive to transformation by LMP2A (ERK) while BCR-ABL1 induced cell death and senescence. In agreement with these findings, BCR-ABL1 is a frequent oncogenic driver in B-ALL but never found in B-cell malignancies past the pre-BCR checkpoint. Likewise, the Epstein-Barr virus (EBV)-encoded oncoprotein LMP2A mimicks pre-BCR signaling^11^ and functions as a common oncogenic driver in mature B-cell malignancies while EBV^+^ B-ALL is exceedingly rare.

## BCL6 promotes survival at the pre-BCR checkpoint and is essential for RAS-mediated B-cell transformation

Our experiments identified activation of BCL6 at the expense of MYC as a central determinant of the pro-B to pre-B cell transition. Interestingly, both initiation of normal pre-BCR signaling and activation of oncogenic ERK (*NRAS*^G12D^) induce strong upregulation of BCL6 (**Fig. 4d**; **Extended Data Figure 7a**). Genetic experiments based on inducible ablation of *Mapk1* (Erk2) and *Stat5* showed that Erk2 is required for upregulation of Bcl6 while Stat5 functions as a negative regulator. On the other hand, Bcl6 is absolutely essential for malignant B-cell transformation by *NRAS*^G12D^. Cre-mediated deletion of *Bcl6* impaired colony formation of *Bcl6*^fl/fl^ *NRAS*^G12D^ B-ALL cells and induced progressive depletion of B-ALL cells. Even deletion of one allele profoundly reduced colony formation and induced cell death, while inducible overexpression of Bcl6 increased colony numbers (**Extended Data Figure 7d-h**). In a transplant model based on murine *Bcl6*^fl/fl^ *NRAS*^G12D^ B-ALL carrying tamoxifen-inducible Cre, inducible deletion of *Bcl6* subverted leukemogenesis *in vivo*, highlighting the essential role of Bcl6 in Ras-mediated B-cell transformation (**Extended Data Figure 7i**). These results in a murine transplant model for *NRAS*^G12D^ B-ALL identified BCL6 as a previously unrecognized vulnerability. To determine whether patient-derived B-ALL cells carrying ERK-activating lesions likewise depend on BCL6 function, we tested pharmacological inhibition of BCL6-function using a *retro-inverso* peptide inhibitor (RI-BPI^33^) and a small molecule inhibitor (FX1^34^). Interestingly, ERK-driven B-ALL PDX cells were more sensitive to both RI-BPI and FX1 when compared to STAT5-driven B-ALL PDX cells. In addition, RI-BPI and FX1 synergized with Trametinib-induced inhibition of oncogenic ERK-signaling (**Extended Data Figure 7j-k**). In patient-derived STAT5-driven B-ALL, Bcl6 expression was repressed but inducible by pharmacological inhibition of STAT5-signaling (Imatinib, Ruxolitinib, Pimozide; **Extended Data Figure 7**). Likewise, pharmacological activation of ERK-signaling by BCI-215 and inducible oncogenic activation of NRAS^G12D^ strongly induced BCL6-expression, while Trametinib repressed BCL6 (**Extended Data Figure 7l-n**). BCI-215 functions as an allosteric small molecule inhibitor of DUSP6^35^, a central negative feedback regulator of ERK in B-ALL^36^.

## Negative regulation of Stat5 by PTPN6 is essential for oncogenic ERK-signaling in B-ALL

Previous work showed that the inhibitory phosphatase PTPN6 (SHP1) is a central negative regulator of STAT5^37^. Inducible activation of a gain-of-function variant of Stat5a (TetO-Stat5a^CA^) repressed Ptpn6, while inducible ablation of Stat5 (Stat5^fl/fl^ +Cre) increased Ptpn6 mRNA levels (**Extended Data Figure 5c-d**). Compared to STAT5-driven B-ALL, PTPN6 is expressed at higher levels in ERK-driven B-ALL (**Extended Data Figure 5g-i**). Since oncogenic ERK-signaling is directly opposed by STAT5 (**Fig. 2-3**), we hypothesized that PTPN6-mediated negative regulation of STAT5 contributes to oncogenic ERK-signaling. Consistent with this scenario, activation of ERK downstream of oncogenic NRAS^G12D^ not only suppressed STAT5 activity but also induced expression and activation of PTPN6 (**Extended Data Figure 8a**). The ERK-downstream transcription factors, ELK1, JUN, JUNB and CREB1^18^ bound to the *PTPN6* promoter, which is consistent with transcriptional activation of *Ptpn6* by NRAS^G12D^-ERK signaling (**Extended Data Figure 8b-c**). In line with a central role of Ptpn6 as negative regulator of Stat5, Cre-mediated deletion of *Ptpn6* increased Stat5-phosphorylation (Y^694^) by >10-fold (**Extended Data Figure 8d**). These findings corroborate that oncogenic activation of Erk negatively regulates Stat5-activity by transcriptional activation of Ptpn6, which then dephosphorylates Stat5 (**schematic Fig. 4j**). Consistent with a critical function of Ptpn6 in oncogenic Erk-signaling, Cre-mediated deletion of *Ptpn6* compromised colony-forming ability of *NRAS*^G12D^ B-ALL cells and induced rapid cell death (**Extended Data Figure 8e-f**).

## The pre-B-cell linker BLNK functions as an insulator to separate Stat5-Myc and Erk-Bcl6 pathways

Previous work showed that the pre-B-cell linker Blnk directly interacts with Ptpn6 and mediates its phosphatase activity^38^. Since Blnk not only induces transcriptional activation of Ptpn6 (**Extended Data Figure 8c**) but also interacts with Ras proteins to promote Erk-signaling^39^, we examined whether Blnk, in addition to Ptpn6, contributes to oncogenic ERK-signaling. Studying inducible activation of pre-BCR- or oncogenic ERK-signaling (**Extended Data Figure 9a-b**) in *Blnk*^+/+^ and *Blnk*^-/-^ B-cell precursors revealed that Blnk was essential for activation of Erk downstream of the pre-BCR or oncogenic NRAS^G12D^. In the absence of *Blnk*, inducible pre-BCR- or NRAS^G12D^-signaling failed to activate Erk and to suppress Stat5-phosphorylation. Likewise, Blnk was required for upregulation of Ptpn6 in response to BCR- or oncogenic Erk-signaling (**Extended Data Figure 9a-b**). Mimicking pre-BCR-downstream signals, oncogenic NRAS^G12D^ induced the transition from Stat5^+^ Myc^+^ pro-B to Erk^+^ Bcl6^+^ B-cells (**Fig. 4a-b**). These results show that both pre-BCR- and NRAS^G12D^-signaling relied on Blnk to successfully complete this transition.

## BLNK enforces convergence of oncogenic signaling on one single pathway

Our gain-of-function experiments showed that concurrent activity of both Stat5^+^ and Erk^+^ pathways (double-positive cells) compromised competitive fitness of leukemia clones and induced senescence and cell death (**Extended Data 3d-i**), unless the conflict was resolved and clones converged on one single pathway (single-positive cells; **Fig. 2e-f**). To examine whether Blnk is required for pathway convergence, oncogenic STAT5- (STAT5a^CA^ or BCR-ABL1) and ERK (NRAS^G12D^) were concurrently activated in murine *Blnk*^+/+^ and *Blnk*^-/-^ B-cell precursors. In the presence of *Blnk*, concurrent activation of Stat5-Myc (GFP^+^) and Erk-Bcl6 (Orange^+^) pathways largely resulted in pathway convergence and outgrowth of Erk-Bcl6 single-positive clones. Compared to 7% in *Blnk*^+/+^ B-cell precursors, the fraction of GFP^+^ Orange^+^ double-positive cells increased to up to 78% in *Blnk*^-/-^ B-cell precursors (**Extended Data Figure 9c**), indicating that Blnk is essential for convergence of oncogenic signaling on one principal pathway (e.g. Erk-Bcl6) and suppression of secondary divergent pathways (e.g. Stat5-Myc; **Extended Data Figure 9d**). Since inducible activation of Stat5a^CA^ suppressed Blnk, while genetic ablation and dominant-negative Stat5 increased Blnk mRNA and protein levels (**Extended Data 5c-e**), we tested whether Blnk-function differentially affects oncogenic Stat5- and Erk-signaling. In the presence of functional Blnk, doxycycline-inducible expression of Stat5a^CA^ failed to transform and *Blnk*^+/+^ B-cell precursors rapidly underwent cell death. In contrast, inducible activation of Stat5a^CA^ in *Blnk*^-/-^ B-cell precursors induced rapid malignant transformation, in agreement with previous findings that Stat5b-CA cooperates with defects in *Blnk* to promote B-ALL^7^. As expected, doxycycline-inducible expression of NRAS^G12D^ caused malignant transformation of *Blnk*^+/+^ B-cell precursors. In the absence of *Blnk*, however, inducible activation of NRAS^G12D^ only transiently accelerated proliferation of B-cell precursors. After 5 days, NRAS^G12D^-transduced *Blnk*^-/-^ B-cell precursors ceased to proliferate *in vitro* and viability progressively declined (**Extended Data Figure 9e-h**). Colony formation assays revealed similar differences between oncogenic Stat5- and Erk-signaling relative to Blnk-function. Ablation of *Blnk* substantially increased the number of colonies after Stat5a^CA^-induction, while colony numbers were decreased in *Blnk*^-/-^ B-cell precursors when Erk was activated downstream of NRAS^G12D^ (**Extended Data Figure 9i-j**). To determine whether BLNK differentially affects oncogenic Stat5- and Erk-signaling also in patient-derived B-ALL cells, we studied a matched B-ALL pair from the same patient at the time of diagnosis (LAX7) and after relapse (LAX7R). LAX7 included a dominant STAT5 clone (*IL7R*^SI246S^) clone whereas LAX7R was driven by a dominant ERK (*KRAS*^G12V^) clone (**Fig. 5a-b**). To validate observations from *Blnk*^-/-^ B-ALL mouse models based on patient-derived B-ALL cells, we used Cas9 ribonucleoproteins (RNPs) and guide RNAs targeting *BLNK* for genetic deletion in LAX7 and LAX7R cells. In agreement with findings in murine B-ALL cells, deletion of *BLNK* provided a strong competitive advantage to STAT5-driven LAX7 (diagnosis) but induced cell death in ERK-driven LAX7R (relapse) B-ALL cells (**Extended Data Figure 9k-m**).

**Figure 5:**
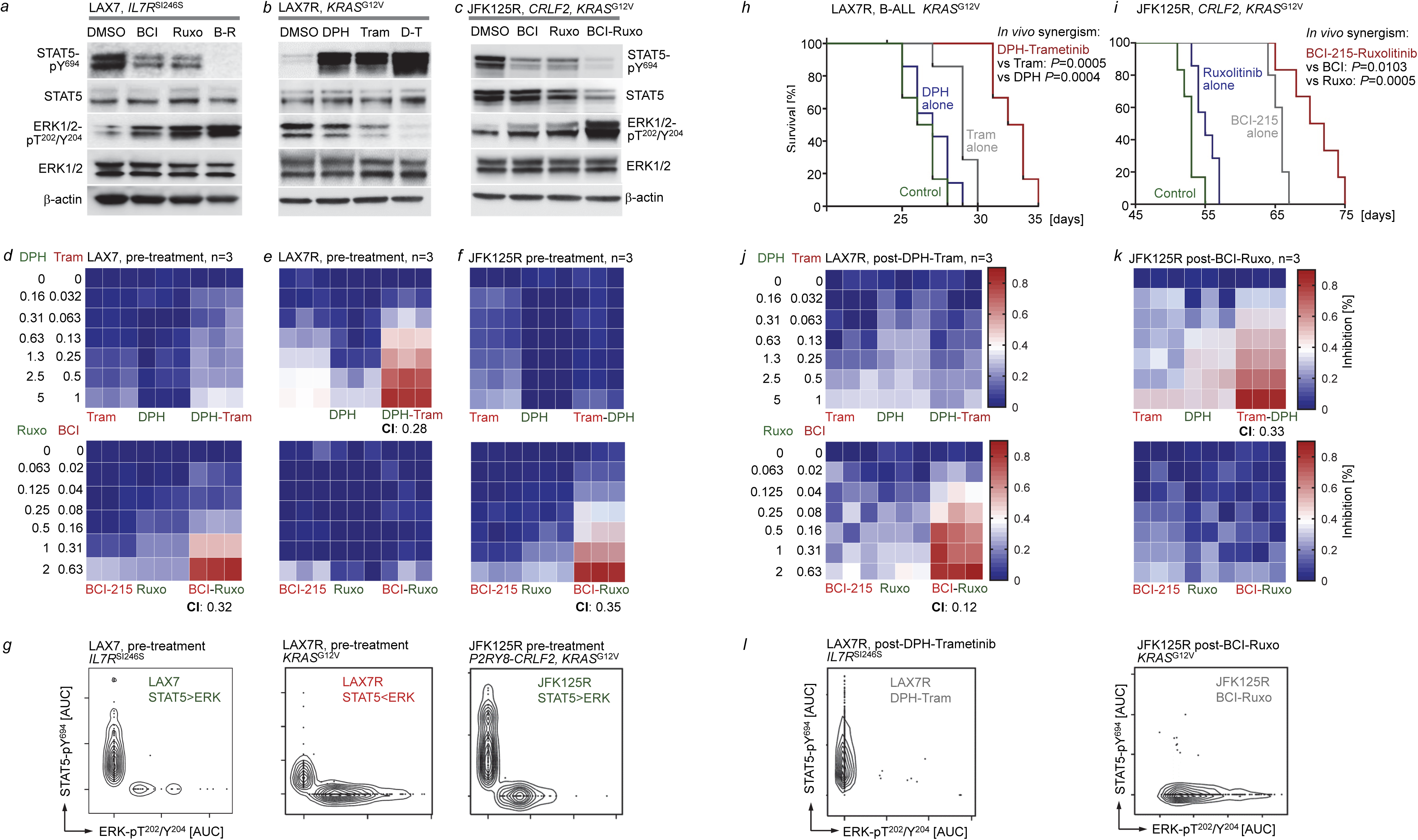
Pharmacological reactivation of suppressed divergent pathways as therapeutic strategy in B-ALL. Patient-derived B-ALL cells (STAT5-driven, LAX7) were treated with vehicle control, BCI-215 (1 µM), ruxolitinib (500 nM), or a combination of BCI-215 and ruxolitinib. Western blotting was performed to measure levels of phospho-STAT5-Y^694^, STAT5, phospho-ERK1/2-T^202^/Y^204^ and ERK1/2. (**b**) Patient-derived B-ALL cells (ERK-driven, LAX7R) were treated with vehicle control, DPH (1 µM), trametinib (500 nM), or a combination of DPH and trametinib. Levels of phospho-STAT5-Y^694^, STAT5, phospho-ERK1/2-T^202^/Y^204^ and ERK1/2 were assessed. (**c**) Patient-derived B-ALL cells (STAT5-driven, JFK125R) were treated and studied by Western blot analysis as described in (**a**). Patient-derived B-ALL cells, LAX7 (**d**), LAX7R (**e**) and JFK125R (**f**) were treated with increasing concentrations of BCI-215, ruxolitinib, a combination of both, DPH, trametinib, or a combination of both. Percentage growth inhibition at each concentration is shown as heatmaps (n=3). CI values were calculated to determine synergy for treatment combinations. (**g**) Single-cell phosphoprotein analyses for phospho-STAT5-Y^694^ and phospho-ERK-T^202^/Y^204^ were performed for patient-derived B-ALL cells LAX7, LAX7R and JFK125R prior to in vivo treatment. (**h-i**) Patient-derived LAX7R and JFK125R B-ALL cells were injected into sublethally irradiated (2Gy) NSG mice. Recipient mice injected with LAX7R were treated 6 times a week for 4 weeks with 2 mg/kg DPH, 0.5 mg/kg trametinib or a combination of both (**h**). Recipient mice injected with JFK125R were treated 5 times a week for 4 weeks with 2 mg/kg BCI-215, 30 mg/kg ruxolitinib or a combination of both (**i**). Mice were euthanized when they showed signs of overt leukemia (hunched back, weight loss and inability to move). Kaplan-Meier survival analyses (Mantel-Cox log-rank test) are shown. To assess additive vs synergistic activity of single vs combination treatments *in vivo*, the Bliss independence model was adapted to survival analysis. With this approach, treatments are Bliss “independent” if the fraction of cells surviving combination therapy equals the product of fractions that survive the individual treatments. A Weibull distribution {exp[−(t/β)α]} was fitted to survival data for each condition, and distributions of survival benefits (treated survival time − untreated survival time) were computed for each treatment. Survival benefits of drug-1 and drug-2 were summed and added to the untreated survival distribution to compose a “sum of benefits” survival distribution. Confidence intervals were based on 10,000 such simulations, in which each treatment’s survival distribution was a likelihood-weighted sample of Weibull parameters (α, β) based on the relative likelihood of having made the survival observations O = (t1, t2,…ti) given those parameters: L(O | α, β) = Πi{exp[−(ti/β)α] × tiα−1 × α × β−α}. The *P* value for synergy between drug-1 and drug-2 was the probability to draw data from the sum of benefits model ensembles with median survival duration equal to or greater than was experimentally observed for drug-1 and drug-2. While combination treatments prolonged overall survival, transplant recipient mice ultimately developed overt leukemia. B-ALL that developed in mice with LAX7R (after treatment with DPH-trametinib; **j**) and in mice with JFK125R (after treatment with BCI-215-ruxolitinib; **k**) were isolated from bone marrow, suspended in cell culture medium and treated with increasing concentrations of BCI-215, ruxolitinib, a combination of both, DPH, trametinib, or a combination of both. Percentage growth inhibition at each concentration is shown as heatmaps (n=3). CI values were calculated to determine synergy for treatment combinations. To elucidate the clonal composition of LAX7R and JFK125R B-ALL post-treatment, single-cell phosphoprotein analyses for phospho-STAT5-Y^694^ and phospho-ERK-T^202^/Y^204^ were performed (**l**).

## Rare occurrence of biclonal disease with distinct STAT5- and ERK-driven B-ALL clones

Single-cell phosphoprotein analyses for STAT5 (Y^694^) and ERK (T^202^/Y^204^) showed a dramatic shift from a dominant Stat5^+^ Erk^-^ clone in LAX7 to a dominant Stat5^-^ Erk^+^ clone in LAX7R at the time of relapse (**Fig. 5**). Few Stat5^-^ Erk^+^ cells were detected in the diagnostic LAX7 sample, that likely gave rise to the dominant clone at the time of relapse (LAX7R), suggesting biclonal disease at the time of diagnosis (**Figure 5g**). Biclonal disease likely represents a rare event: In our analysis of genetic STAT5- and ERK-driver lesions in 1,148 B-ALL samples, we only found evidence for biclonal disease in ∼3% of cases (**Fig. 1a**). In our collection of 59 B-ALL PDX, based on exome sequencing, Western blot and single-cell phosphoprotein analyses, we found 5 cases (8.5%) with two distinct STAT5- and ERK-driven clones (**Fig. 1i, Supplementary Table 4**). The divergent genetic and phenotypic changes between the diagnostic (LAX7R and relapse (LAX7R) sample were mirrored by contrasting drug-responses. LAX7 and LAX7R samples were selectively sensitive to inhibitors of oncogenic JAK-STAT5 (ruxolitinib) and MAP2K1-ERK (trametinib), respectively (**Extended Data Figure 10c**). Interestingly, pharmacological inhibition of the principal oncogenic pathway in LAX7 (STAT5) and LAX7R (ERK) resulted in reactivation of the divergent suppressed pathway (**Extended Data Figure 10d**). The finding that STAT5- and ERK-activating lesions are mutually exclusive suggests the existence of a pool of leukemia-initiating cells that harbored neither STAT5-nor ERK-activating lesions. Subsequent acquisition of STAT5- and ERK-activating lesions in distinct clones would give rise to biclonal disease.

## Pharmacological reactivation of suppressed divergent pathways as therapeutic strategy in human B-ALL

The current paradigm of targeted therapy in cancer is mainly based on direct suppression of the principal oncogenic driver. For instance, inhibitors of the BCR-ABL1 and JAK2 tyrosine kinases have been developed to target oncogenic STAT5 signaling^5,40^ as well as MAP2K1 and BRAF inhibitors to target oncogenic ERK^9,10^. Direct inhibition of a principal oncogenic driver typically results in initial remission followed by development of subsequent drug-resistance^41^. As a strategy to prevent drug-resistance and relapse, we explored an alternative approach based on pharmacological reactivation of suppressed pathways that operate in divergent directions relative to the principal oncogenic driver. For proof-of-concept studies, we focused on pharmacological reactivation of suppressed ERK- and STAT5-signaling by two small molecule compounds. BCI-215 functions as an allosteric inhibitor of DUSP6^35^, a central negative feedback regulator of ERK in B-ALL^36^. The small molecule DPH functions as ABL1 kinase activator upstream of STAT5 by engaging the ABL1 myristoyl binding site to prevent its autoinhibitory conformation^42^. Treatment of patient-derived STAT5-driven B-ALL cells (PDX2; *BCR-ABL1*) with BCI-215 rapidly reactivated suppressed ERK, which was paralleled by increased phosphorylation of PTPN6 and dephosphorylation of its substrate STAT5 (**Extended Data Figure 10e-f**). In two additional patient-derived STAT5-driven B-ALL PDX (LAX7, JFK125), treatment with BCI-215 had similar effects and reactivated the repressed ERK-pathway at the expense of STAT5-phosphorylation (**Fig. 5a-c**). These results suggest that pharmacological reactivation of divergent ERK-signaling (e.g. by BCI-215) can be leveraged for suppression of STAT5 as the principal oncogenic driver. In a converse experiment, we tested pharmacological reactivation of divergent STAT5 (by DPH) in ERK-driven B-ALL cells (LAX7R; KRAS^G12V^). As expected, treatment of LAX7R cells with the STAT5-agonist DPH resulted in reactivation of suppressed STAT5-signaling. Strikingly, in addition to reactivation of divergent STAT5-signaling, DPH induced suppression of ERK-phosphorylation, the principal oncogenic driver downstream of KRAS^G12V^ (**Fig. 5b**). Pharmacological reactivation of suppressed divergent pathways is of particular interest in cases where the principal pathway has acquired on-target drug-resistance, e.g. mutations in BCR-ABL1 or MAP2K1^43^ that confer resistance to ABL1 and MEK kinase inhibitors. In addition, this strategy may be useful when direct inhibitors of the principal oncogenic pathway are not or not well established (e.g. NRAS, KRAS).

## Pharmacological reactivation of divergent pathways synergizes with direct inhibition of the principal driver

We next tested whether pharmacological reactivation of divergent pathways can amplify the impact of direct inhibition of the principal oncogenic driver. For instance, BCI-215 reactivated divergent ERK-signaling and suppressed STAT5 to a similar degree as direct STAT5-pathway inhibition by Ruxolitinib (**Fig. 5a, c**). While BCI-215 or Ruxolitinib alone achieved a 3-to 4.5-fold reduction of STAT5-phosphorylation, a combination of both suppressed levels of STAT5-pY^694^ to near or below detection limits (**Figure 5a,c**). Consistent with biochemical inhibition of STAT5, single-agent treatment with BCI-215 or Ruxolitinib alone only achieved minor responses, whereas a combination of both had a profound synergistic effect in eliminating B-ALL cells (CI: 0.32, **Fig. 5d** and CI:0.35, **Fig. 5f**). In ERK-driven (*KRAS*^G12V^) B-ALL cells (**Fig. 5b**), Trametinib directly suppressed the principal pathway, while DPH reactivated divergent STAT5-signaling. As single-agents, DPH and Trametinib partially suppressed ERK-phosphorylation (**Fig. 5b**) and had minor effects on cell viability. Mirroring complete suppression of ERK-phosphorylation, combinations of DPH and trametinib had profound cytotoxic effects on B-ALL cells (CI:0.28; **Fig. 5e**). To validate efficacy and feasibility of divergent pathway reactivation in an *in vivo* transplant setting, we injected immunodeficient NSG mice with LAX7R and JFK125R B-ALL PDX cells. In both cases, STAT5-driven primary B-ALL had developed drug-resistance and relapsed with an additional *KRAS*-driven clone (ERK). In LAX7R, the ERK-clone had developed from a small population at the time of diagnosis and become dominant at the time of relapse (**Fig. 5g**). In JFK125R, the additional clone at the time of relapse was still minor (**Fig. 5g**). Treatment of mice bearing LAX7R with a combination of DPH and Trametinib substantially extended overall survival of mice bearing ERK-driven B-ALL (*KRAS*^G12V^), compared to mice that were treated with either compound alone (**Fig. 5h**). DPH in combination with Trametinib achieved synergy based on a significantly superior survival (*P*=0.0001) compared with that expected from additivity of DPH and Trametinib applied alone. Since the STAT5-driven clone in JFK125R at the time of relapse was still dominant (**Fig. 5g**), mice bearing JFK125R were treated with combinations of BCI-215 and Ruxolitinib. Compared to single-agent treatment, combinations of BCI-215 and Ruxolitinib significantly prolonged overall survival of transplant recipients. BCI-215 in combination with Ruxolitinib produced a duration of survival that was significantly longer than an “additivity” model based on the sum of survival benefits of BCI-215 and ruxolitinib applied separately, with *in vivo* synergy (*P*=0.0009).

Compared to inhibitors of the principal oncogenic driver alone, combinations with divergent pathway activation substantially deepened drug-responses and prolonged overall survival of transplant recipient mice. However, eventually all mice died from leukemia in both models, LAX7R (dominant ERK) and JFK125R (dominant STAT5). Our genetic (**Fig. 1a, Supplementary Table 4**) and biochemical (**Fig. 1i**) studies revealed that >90% of B-ALL cases comprise only a single population that is either STAT5- or ERK-driven. LAX7R and JFK125R were not only refractory but also harbored both STAT5- and ERK-driven clones. Hence, we examined how treatment with DPH-Trametinib and BCI-Ruxolitinib combinations affected the clonal composition of B-ALL that ultimately developed fatal disease. To this end, we compared STAT5- and ERK-phosphorylation by single-cell analysis of LAX7R and JFK125R before (**Fig. 5g**, pre-treatment) and after treatment with DPH-Trametinib and BCI-Ruxolitinib combinations *in vivo* (**Fig. 5l**, post-treatment). Interestingly, in both B-ALL PDX, combination treatment eradicated the dominant clone, whereas the formerly minor clone gave rise to fatal disease driven by one single population (**Fig. 5l**). We also compared drug-responses pre- and post-treatment *in vivo*. Compared to pre-treatment, fatal leukemia post-treatment was resistant to the drug-combination directed at the major clone but had become highly sensitive to the drug combination directed at the divergent pathway (**Fig. 5j-k**). For both LAX7R and JFK125R, fatal leukemia arose from a minor diverging clone rather than acquired drug-resistance of the major clone. In addition, combination treatments successfully eradicated the previously dominant clone, changing the biclonal disease to a single population driven by STAT5 or by ERK (**Fig. 5l**). In addition, the dramatic changes in drug-sensitivity suggest that alternating treatment regimens with (i) a drug-combination targeting the major clone followed by (ii) a drug-combination targeting the divergent minor clone will be useful for eradication of high-risk bi-clonal disease.

*In vivo* treatment studies not only confirmed synergistic effects of BCI-215-Ruxolitinib and DPH-Trametinib combinations but also supported feasibility. Combinations of principal pathway inhibition with divergent pathway reactivation had strong anti-leukemia effects in *vitro* and in *vivo* but did not cause any significant weight loss or other signs of toxicity in mice. These results indicate that convergence on one principal oncogenic driver represents a critical event during B cell transformation and leukemia-initiation and a previously unrecognized vulnerability. Hence, targeted reactivation of divergent pathways to interfere with the principal oncogenic driver can be leveraged as a novel strategy to disrupt oncogenic signaling and overcome drug-resistance.

## Discussion

Non-transformed B-cell precursors constantly exchange information with their environment and depend on external cues for proliferation and survival that engage multiple divergent pathways (e.g. cytokine receptors, pre-BCR). Multiple cytokine receptors and the pre-BCR mainly signal through one pathway but not in a strictly exclusive manner. Hence, a diverse spectrum of signaling input reflects interactions of normal cells with their environment, while convergence on one central pathway represents a hallmark of a malignant (cancer or leukemia) state. In addition, signals in response to exogenous stimuli of normal growth factor receptors are typically transient. In contrast, genetic lesions and oncogenic mimics of cytokine receptor or pre-BCR signals result in persistent activation, which could explain that the signal output from these oncogenic mimics is more centralized on one single pathway than signal transduction from normal cytokine receptor or pre-BCR signaling. The dependency of normal B-cells on signaling input from a diverse repertoire of surface receptors is in contrast to “self-sufficiency in growth signals”, one of the hallmarks of cancer cells^44^. “Self-sufficiency” reflects that transforming oncogenes often mimic survival and proliferation signals that would emanate from these receptors, expression and activity of which is no longer needed in transformed B-cells. For instance, transforming oncogenes in B-cell leukemia and lymphoma frequently engage cytokine receptor^7^ or (pre-)BCR-downstream^11,45^ signaling pathways. As a result of oncogenic mimicry of these pathways, “self-sufficient” transformed B-cells lose cytokine receptor or BCR expression and function^12^. Beyond the established concept of “self-sufficiency” and the implication that multiple cell surface receptors become dispensable in transformed cells, we here provide evidence that inactivation of divergent pathways that are not aligned with the principal oncogenic represents a critical step during malignant transformation. Tracking early stages of leukemia-initiation, we identified convergence on one principal oncogenic driver and inactivation of diverging pathways as critical events during B cell transformation. Our results support a scenario in which reactivation of divergent and potentially conflicting signaling pathways represents a powerful barrier to malignant transformation. In agreement with deletion of *Mapk1* as a leukemia-initiating event in Stat5-driven B-ALL, a recent genetic screen identified truncation of the ERK-signaling activator *Sos1* as a frequent lesion that may trigger the onset of STAT5-driven B-ALL^27^. Conversely, recent observations in patients treated with JAK1/2-inhibitors exemplified how pharmacological suppression of (divergent) STAT5-signaling could prime dormant B-cell clones to develop B-cell lymphoma driven by ERK as principal oncogenic pathway^28^. In this clinical trial for patients with STAT5-driven (JAK2^V617F^) myeloproliferative neoplasms (MPN), pharmacological suppression of STAT5 by JAK1/2-inhibitors was associated with a 15-fold increased risk of mature B-cell lymphomas. From three of these patients, matched MPN and subsequent B-cell lymphoma samples were available. In all three cases, the overt lymphoma was preceded by an abnormal B-cell clone in the MPN sample, including one case with a *KRAS*^G13D^ mutation^28^. Based on our finding that Ruxolitinib accelerates colony formation of *KRAS*^G12D^-driven B-ALL (**Extended Data 4b-d**), results from genetic screens^27^ and clinical trials^28^, it could be important to study if divergent pathway inhibition in other settings can initiate overt B-cell malignancies from silent preexisting clones.

From a treatment-perspective, targeted reactivation of divergent pathways can interfere with the principal oncogenic driver and may represent a powerful strategy to amplify treatment-responses and prevent drug-resistance. Our results show that this approach is orthogonal to direct inhibition of driver oncogenes (e.g. by TKI) and may represent a previously unrecognized strategy to overcome conventional mechanisms of drug-resistance. It is likely that these pathways interact differently in other cell types. In addition, activation of other pathways (e.g. WNT, NF-κB, Notch, Hippo) may either coalesce into one convergent pathway or interfere and disrupt signaling from the principal driver oncogene. Cross-inhibition and reciprocal feedback control was previously observed in multiple cancer types. For instance, in breast cancer and lung cancer, ERK-signaling results in suppression of oncogenic receptor tyrosine kinases (RTKs)^14,15^. Likewise, PI3K and androgen receptor signaling are engaged in a negative feedback loop^16^ and oncogenic mTOR signaling suppresses alternative growth factor signals from multiple RTKs^17^. In all these cases, inhibition of one pathway results in feedback activation of the other (**Extended Data Figure 10g**). Unlike B-ALL, where reactivation of a divergent pathway suppresses the principal pathway and compounds toxicity, activation of an alternative pathway in solid tumors represents a route for survival and drug resistance. Hence, dual suppression of both competing pathways, e.g. by inhibitor combinations, represents an effective strategy to prevent drug-resistance in solid tumors. Combinations of principal pathway inhibitors with reactivation of divergent pathways has not been explored in solid tumors. While current treatment treatment approaches for drug-resistant cancer are focused on drug-combinations to inhibit multiple pathways. Here, we introduce a scenario that is based on inhibition of the principal pathway combined with reactivation of divergent pathways. To comprehensively identify targets for pharmacological reactivation of divergent (suppressed) pathways, it will be important to elucidate cell type-specific mechanisms of convergence and interference between principal oncogenic drivers and their potential detractors.

## Acknowledgments

We would like to thank Lars Klemm, Franziska Auer, Haytham Khoury and Janet Winchester for their help with technical aspects of some of the experiments and current and former members of the Müschen laboratory for their support and helpful discussions. Research in the Müschen laboratory is funded by the NIH through an NCI Outstanding Investigator Award R35CA197628 (to M.M.), U10CA180827 (to E.P., A.M. and M.M.), R01CA137060, R01CA157644, R01CA172558 and R01CA213138 (to M.M.), a Wellcome Trust Senior Investigator Award (WT-101880) and a Leukemia and Lymphoma Scholar award (to M.M.), the Howard Hughes Medical Institute HHMI-55108547 (to M.M.), the Norman and Sadie Lee Foundation (for Pediatric Cancer, to M.M.), and the Falk Trust through a Falk Medical Research Trust Catalyst Award (to M.M.), the Pediatric Cancer Research Foundation (PCRF), the Cancer Research Institute (CRI) through a Clinic and Laboratory Integration Program (CLIP) grant (to M.M.) and the California Institute for Regenerative Medicine (CIRM) through DISC2-10061. M.M. is a Howard Hughes Medical Institute (HHMI) Faculty Scholar.

## Author contributions

L.N.C., R.C., M. Murakami, C.H., K.K., T.S., S.S., C.H., A.U., M.A.N., Z.C., J.W.L., and K.N.C performed experiments and analyzed data. H.G. performed all biostatistical analyses and performed power calculations for experimental design. P.P. performed analysis of TARGET data. C.W.C. and J.C. provided expertise in gene editing experiments. M.H. provided expertise in bioinformatics analysis. A.V. and D.W. provide critical materials. A.W. provided expertise in cell surface proteomics. S.I. and O.L. provided expertise in leukemia biology. M. Müschen secured funding, developed the concept and wrote the manuscript. All co-authors reviewed and edited the manuscript.

## Declaration of Interests

The authors have no competing interests.

## Figure legends

**Extended Data Figure 1:**
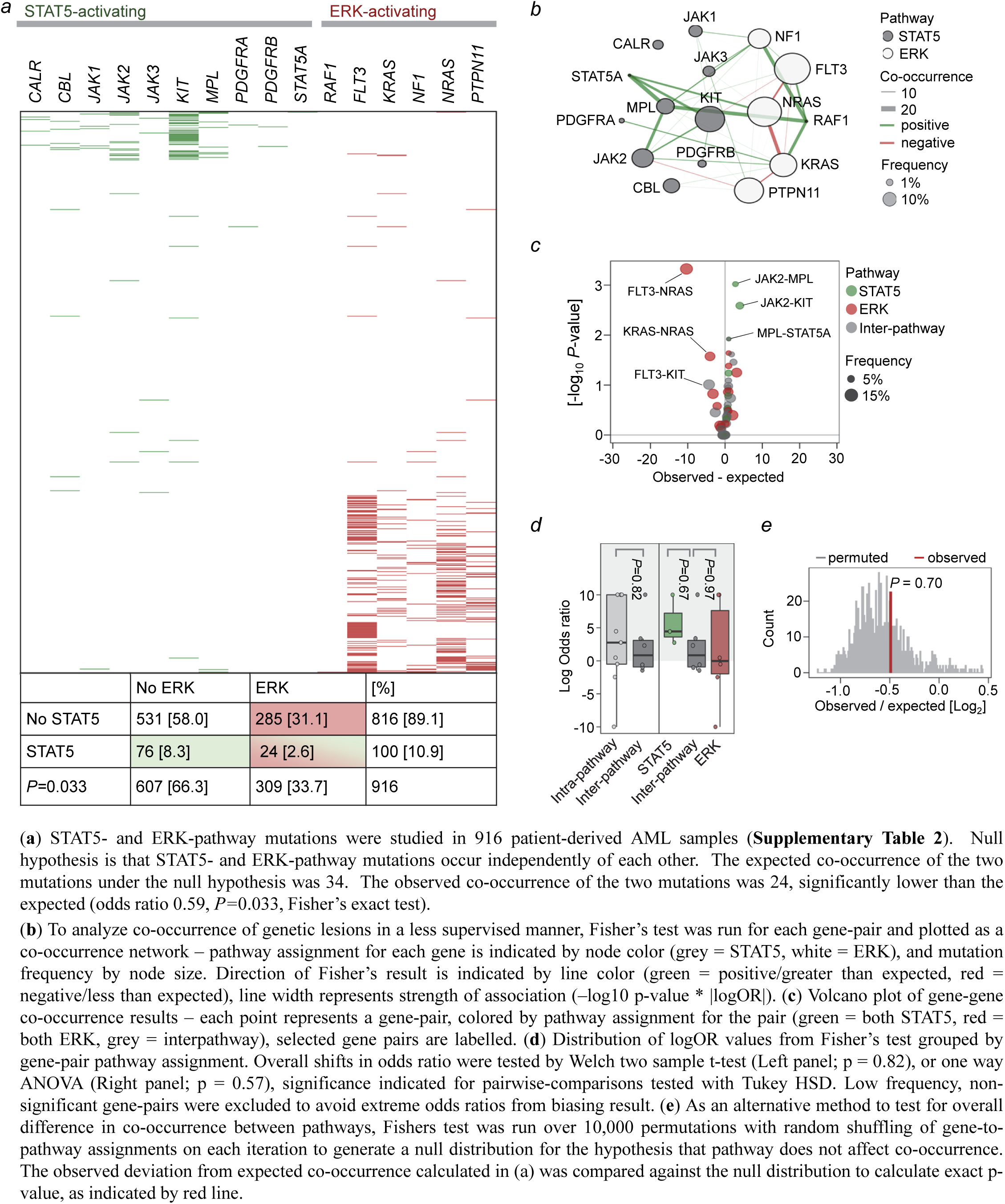
Segregation of STAT5- and ERK-activating mutations in human AML

**Extended Data Figure 2:**
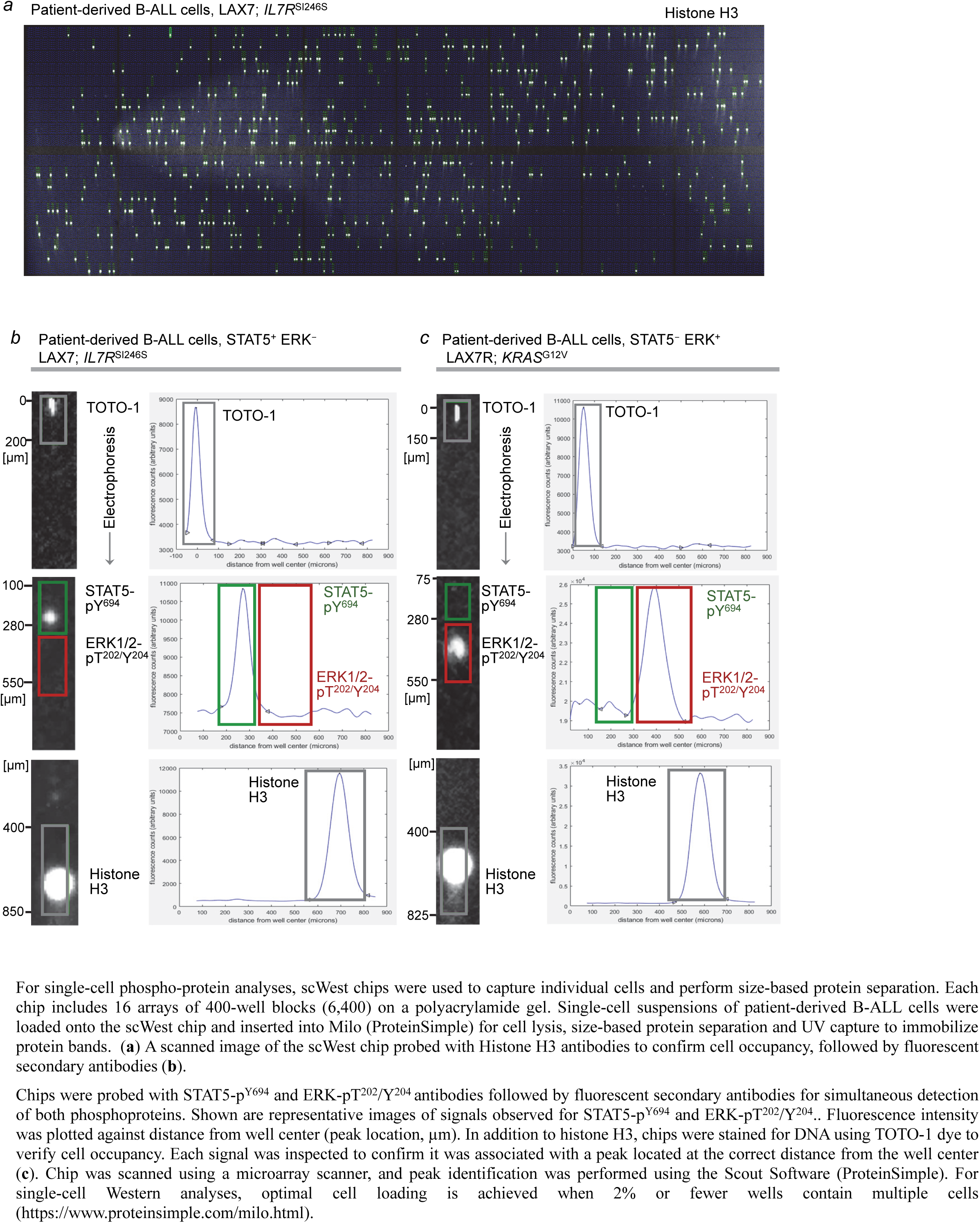
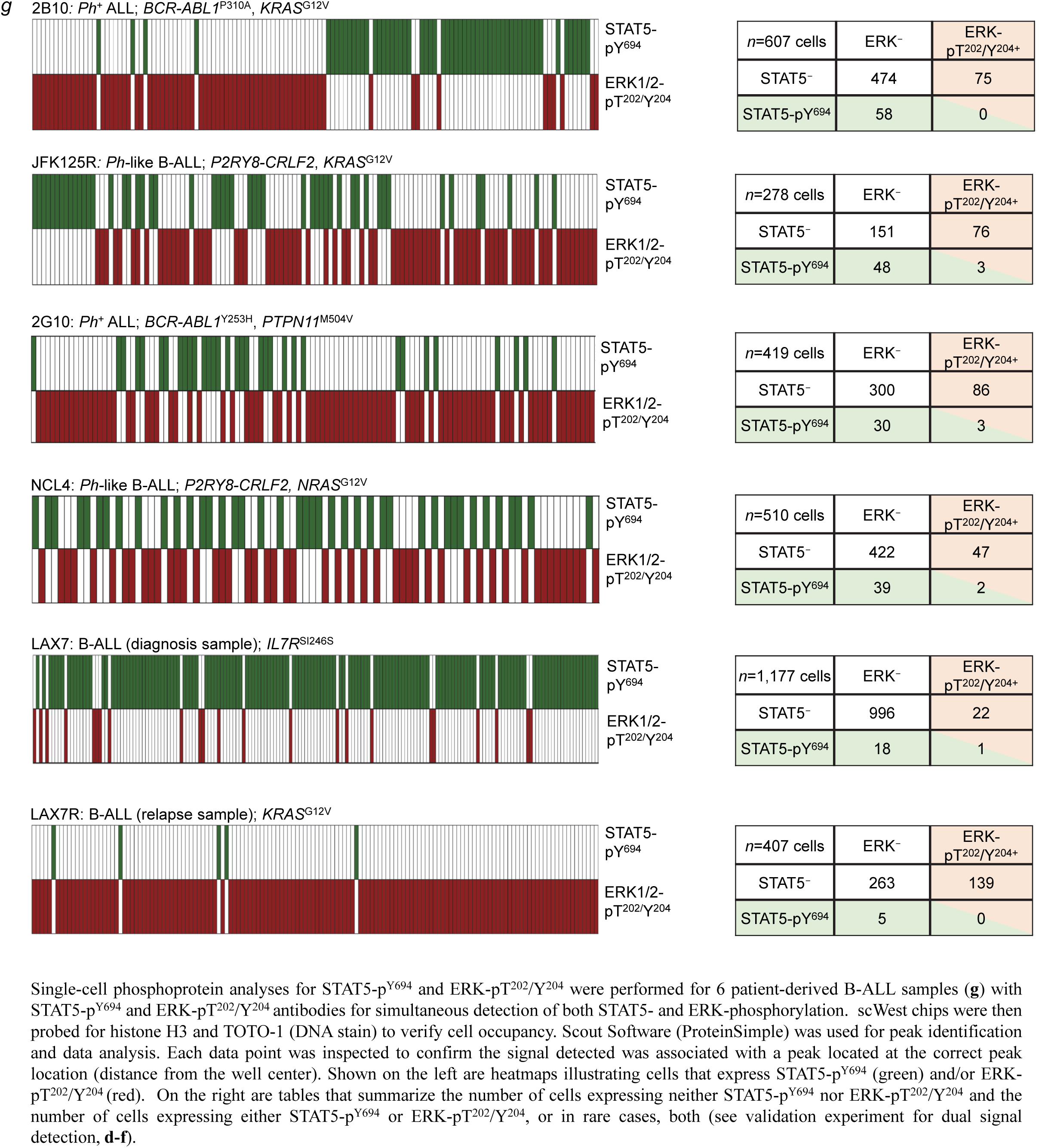
Single-cell phosphoprotein analyses of patient-derived B-ALL samples reveal segregation of STAT5- and ERK-phosphorylation to competing clones

**Extended Data Figure 3:**
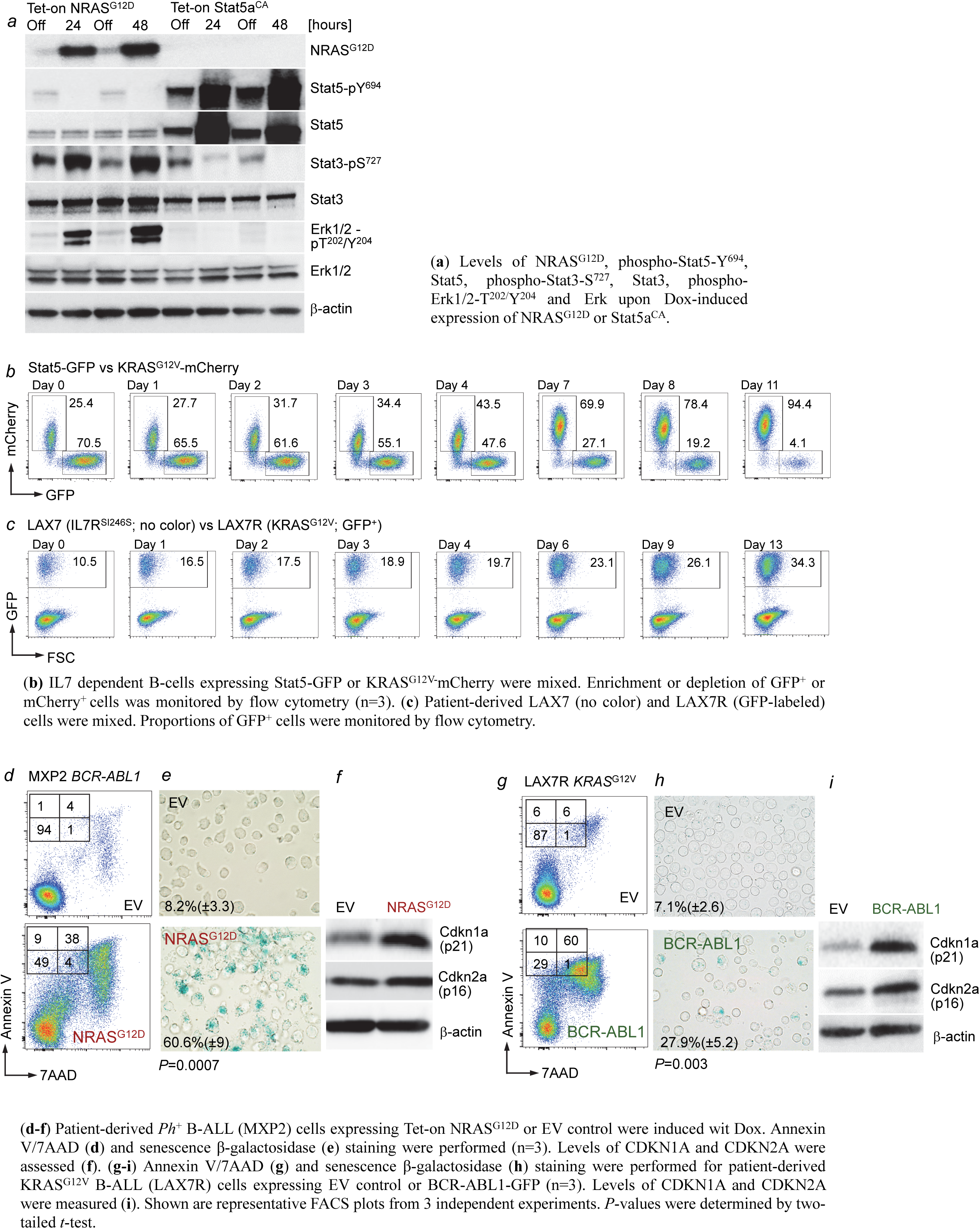
Concurrent activation of STAT5 and ERK signaling induces B-cell senescence and cell death

**Extended Data Figure 4:**
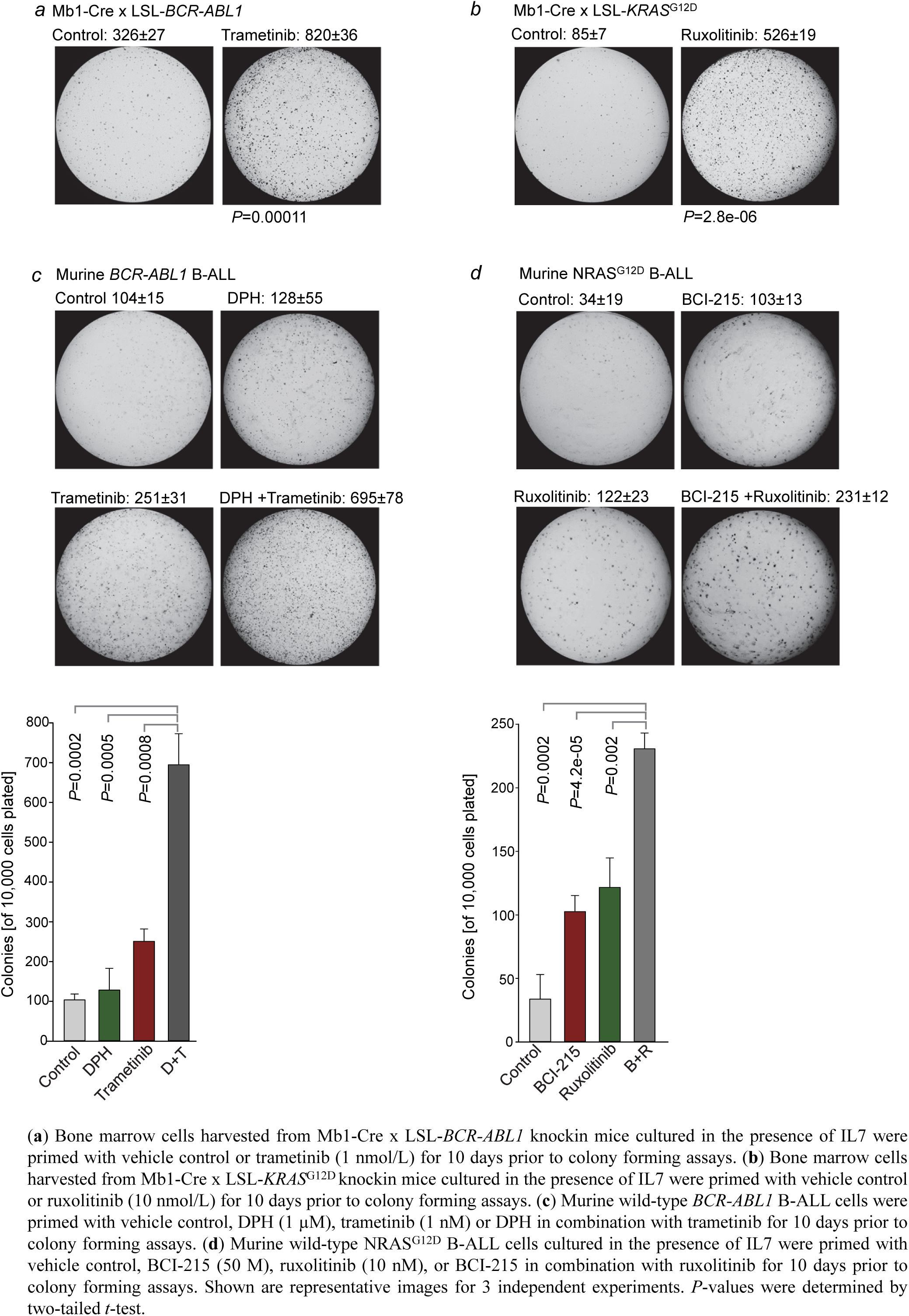
Pharmacological inhibition of divergent pathways facilitates B-leukemogenesis

**Extended Data Figure 5:**
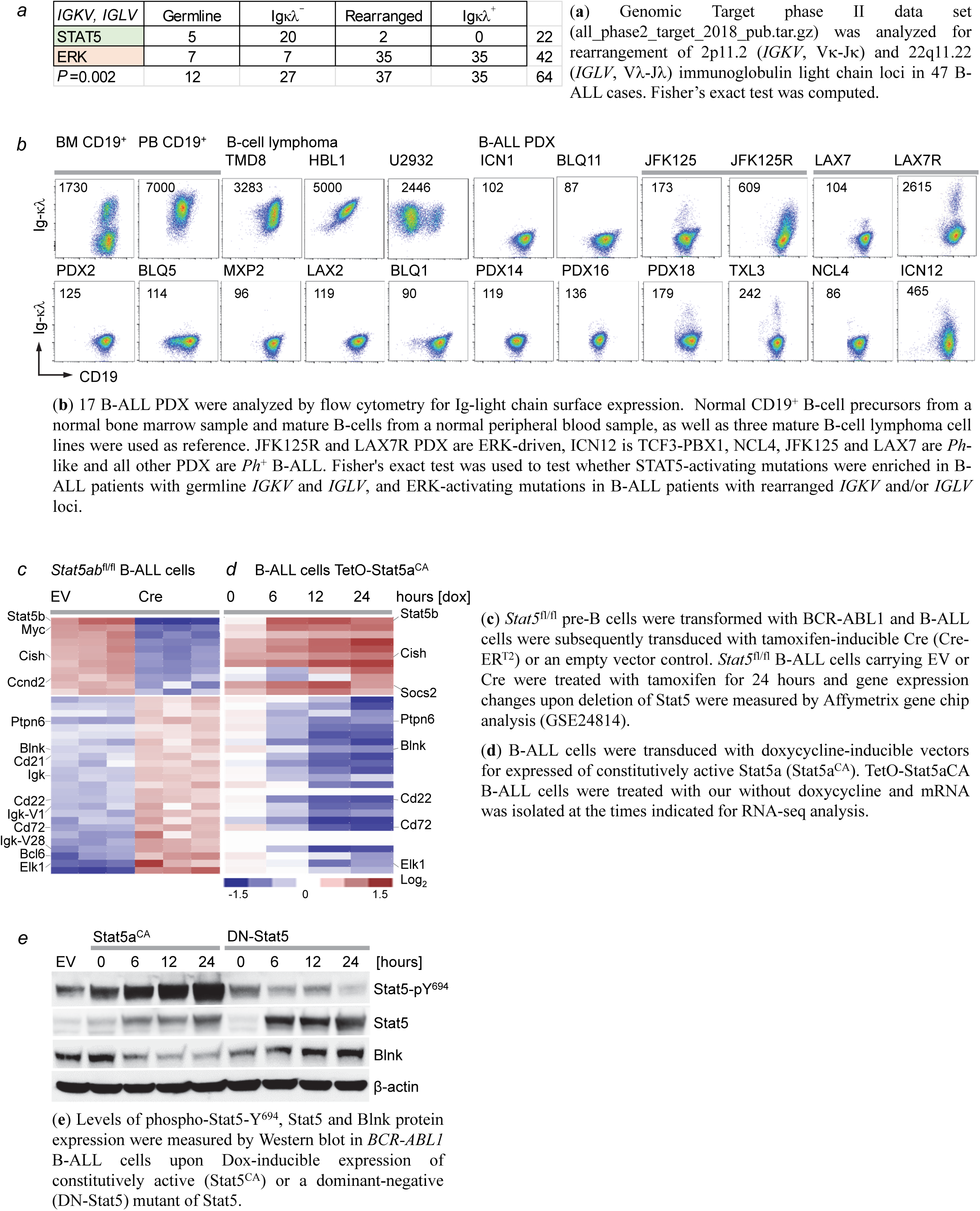
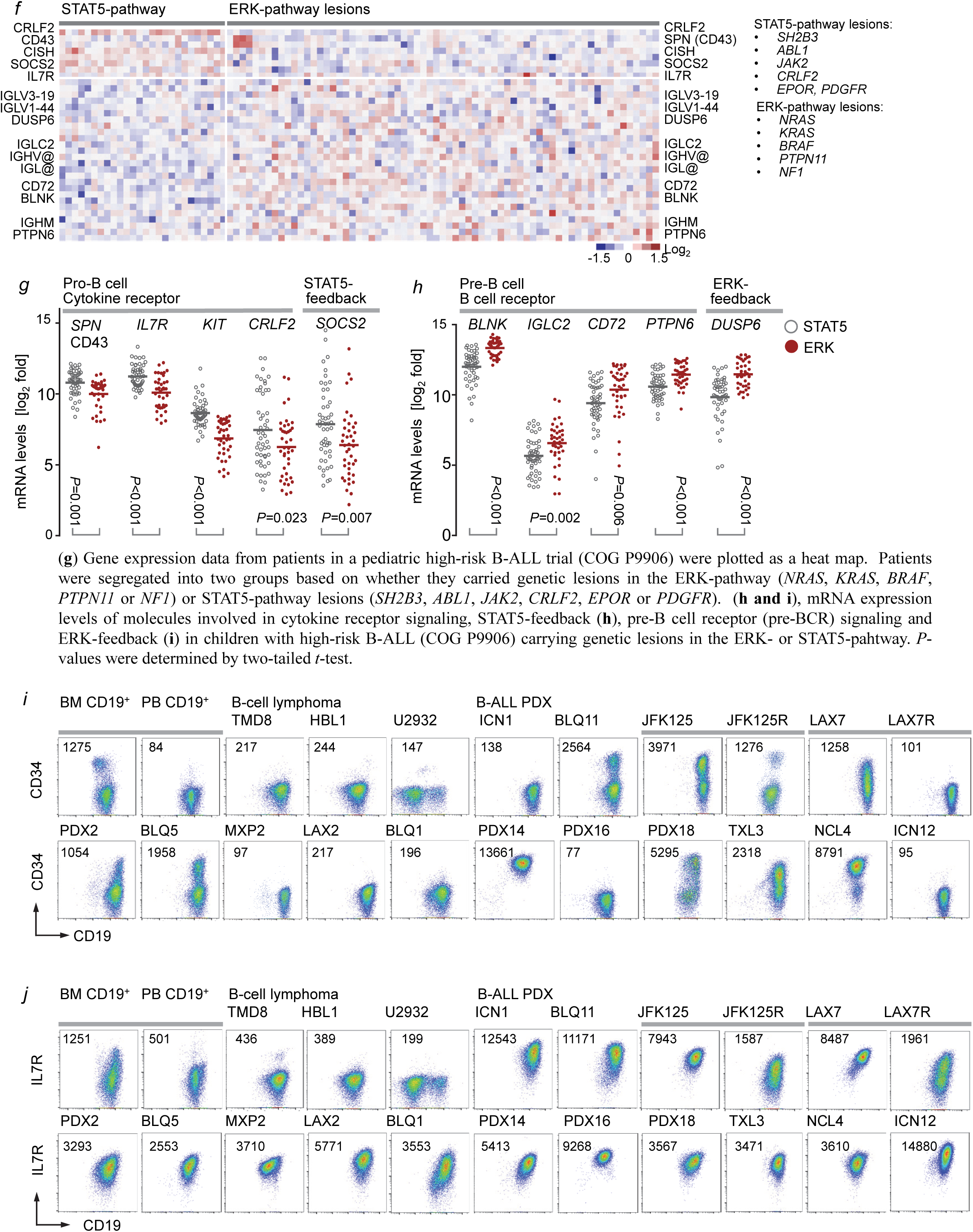
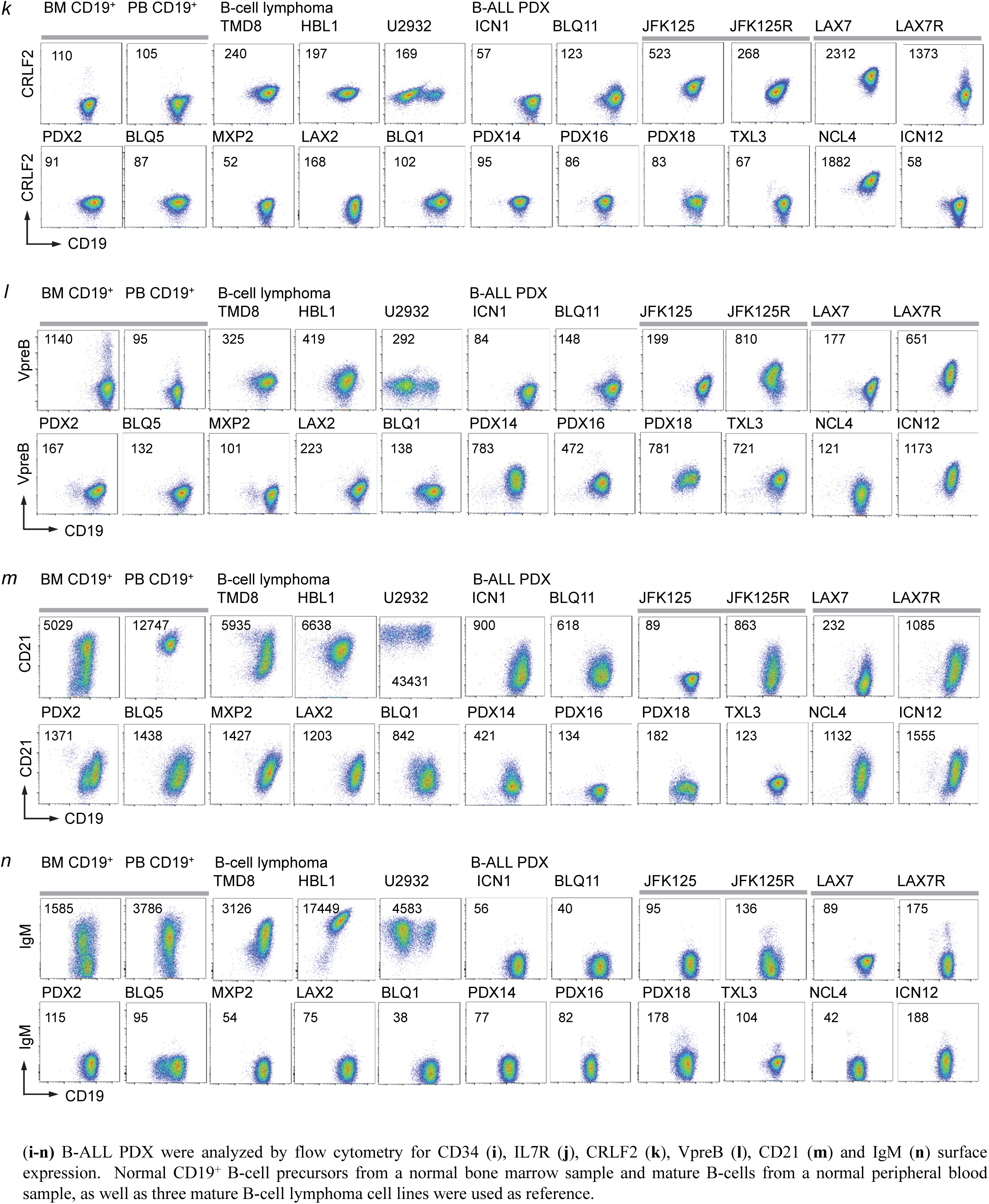
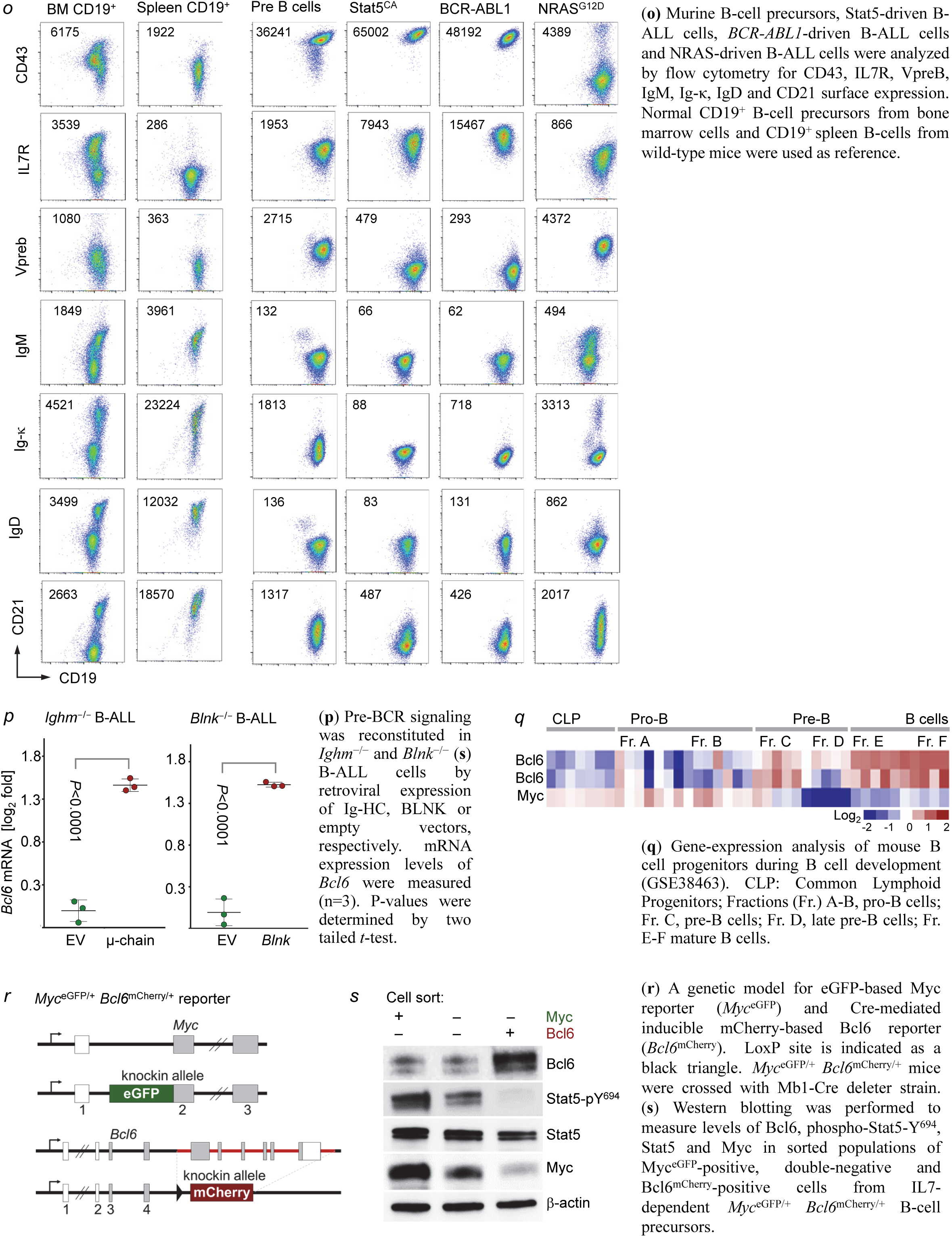
Phenotypic differences between B-ALL cells driven by oncogenic STAT5 or ERK reflect developmental rewiring during early B-cell development

**Extended Data Figure 6:**
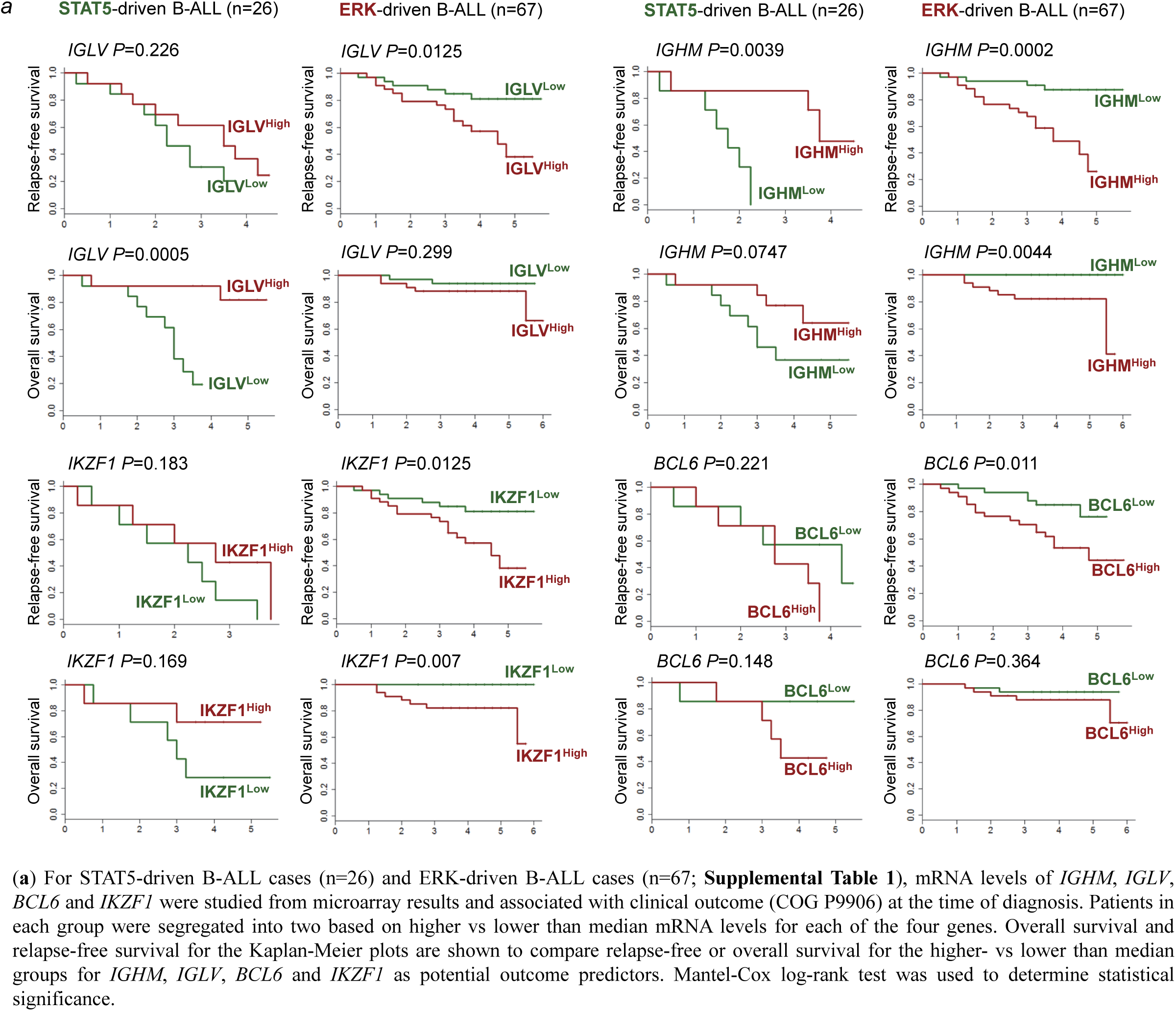
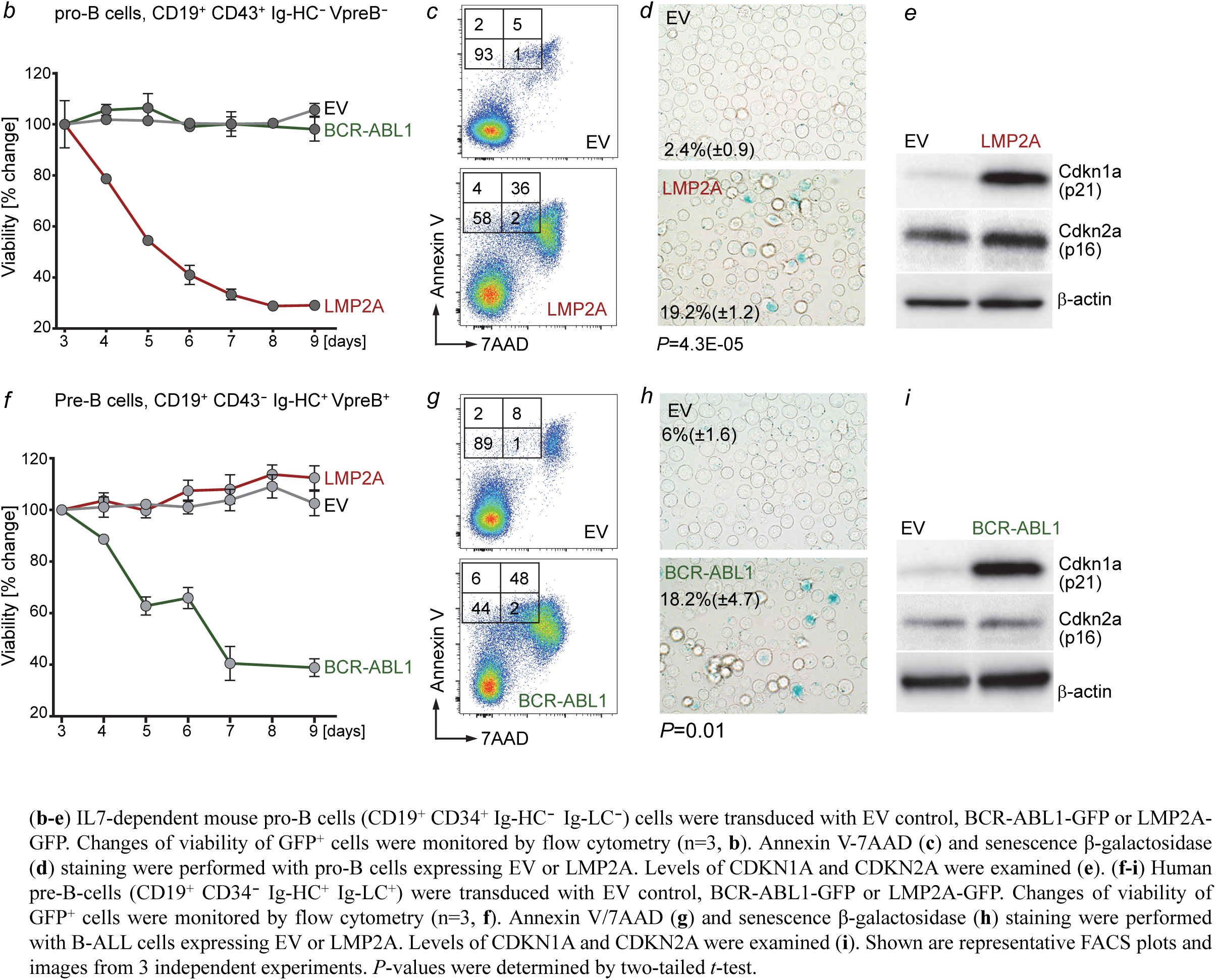
Permissiveness to oncogenic STAT5 or ERK signaling depends on B-cell differentiation stage

**Extended Data Figure 7:**
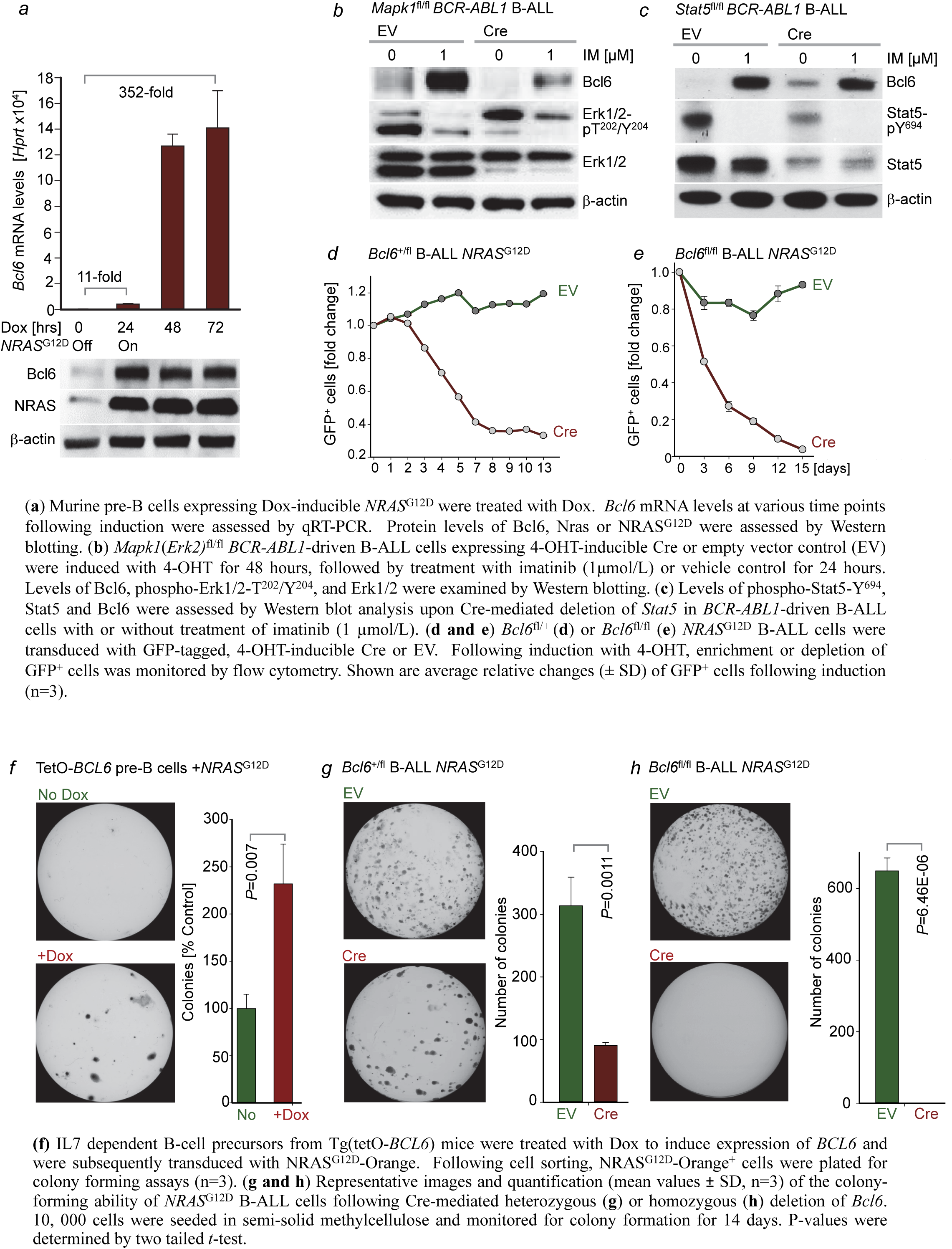
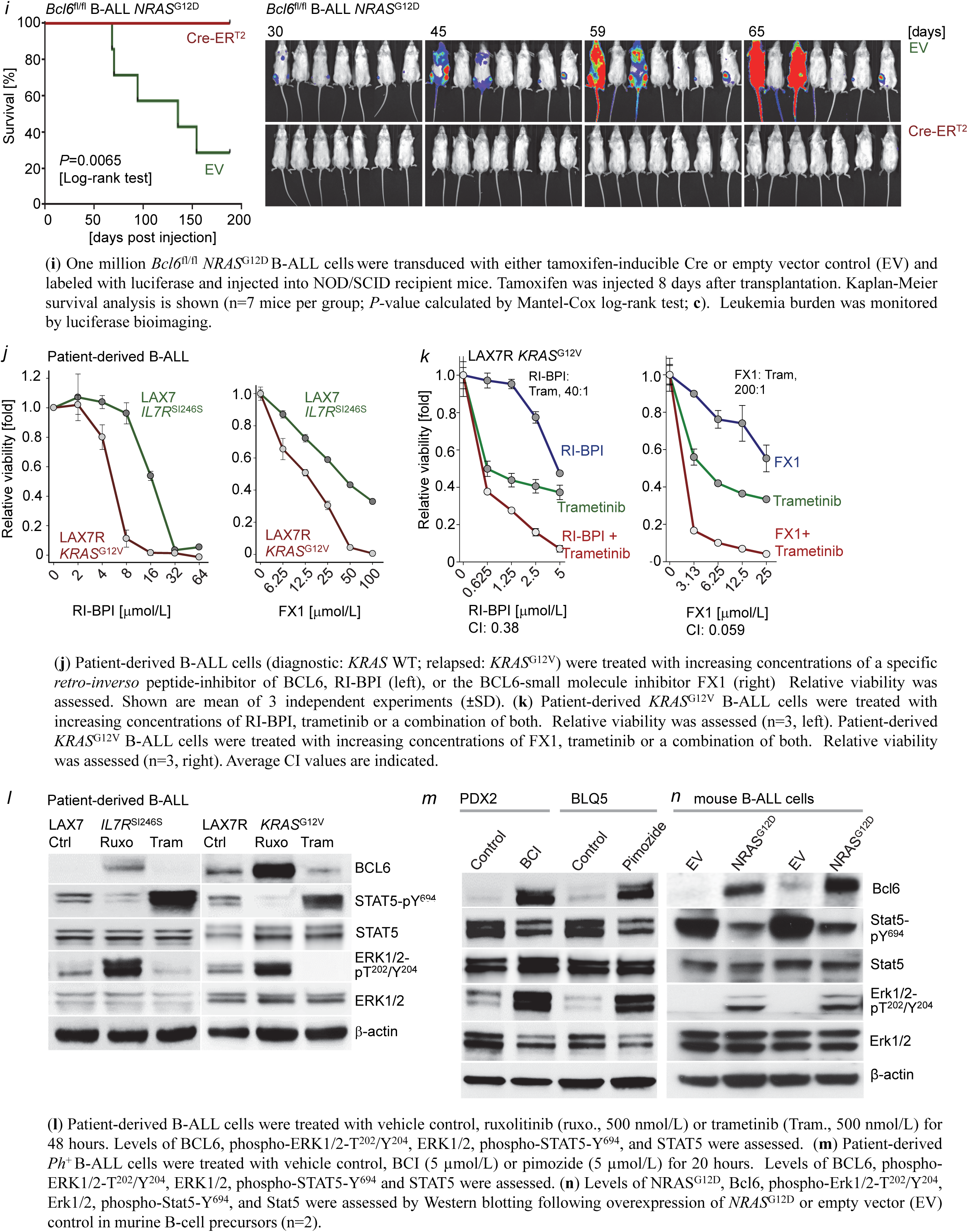
BCL6 is essential for RAS-mediated B-cell transformation

**Extended Data Figure 8:**
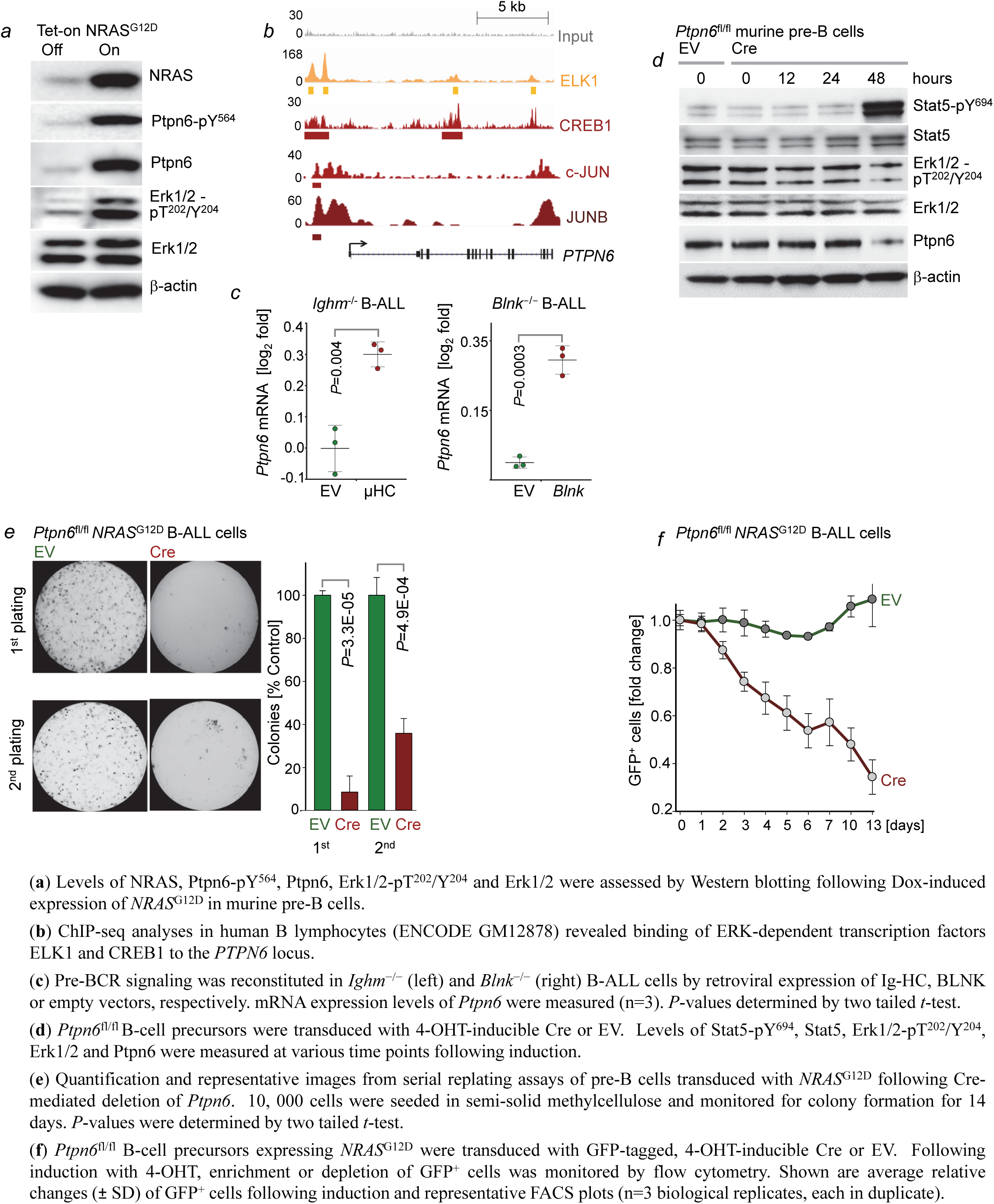
Central role of PTPN6 in enabling oncogenic ERK-signaling

**Extended Data Figure 9:**
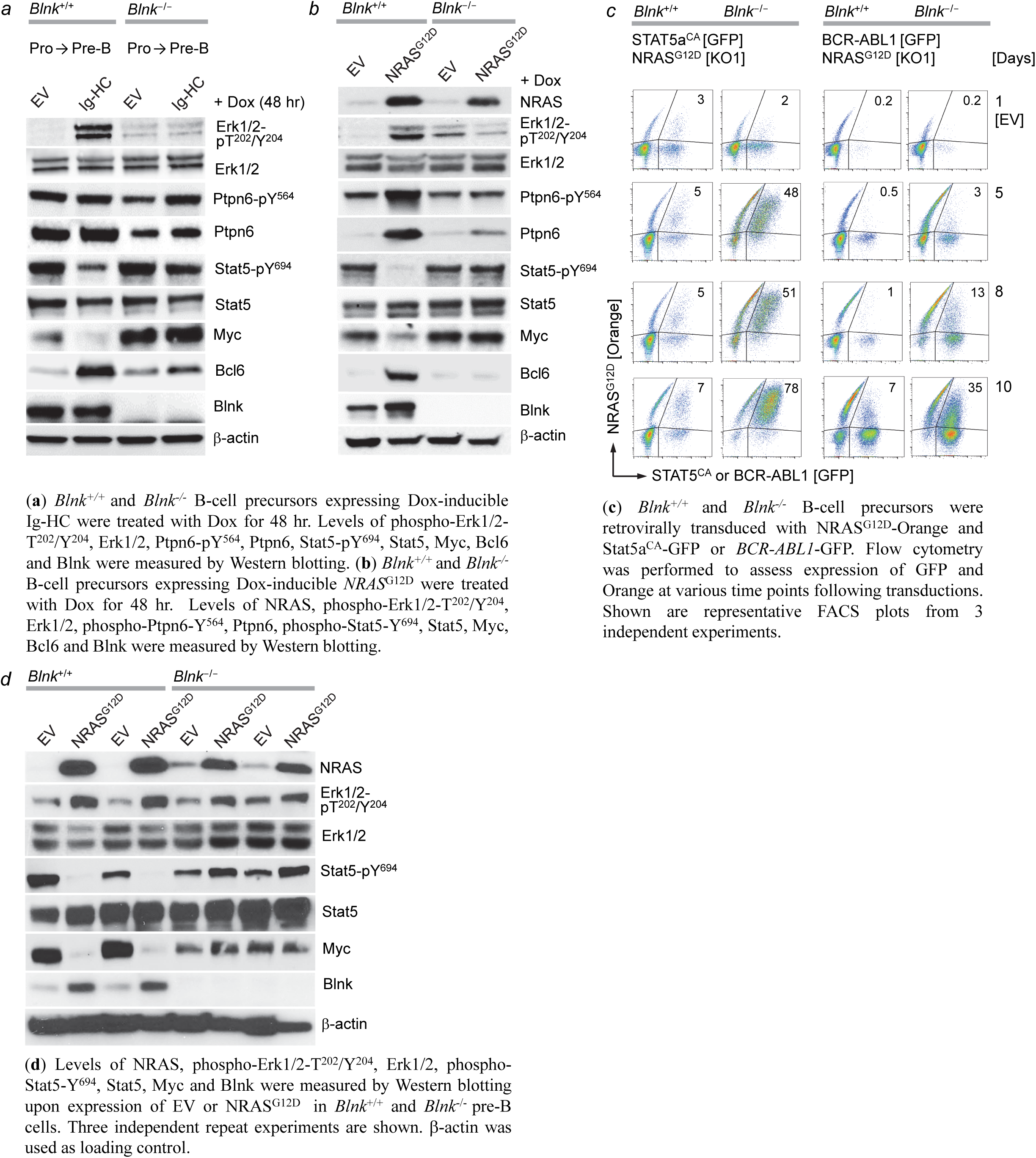
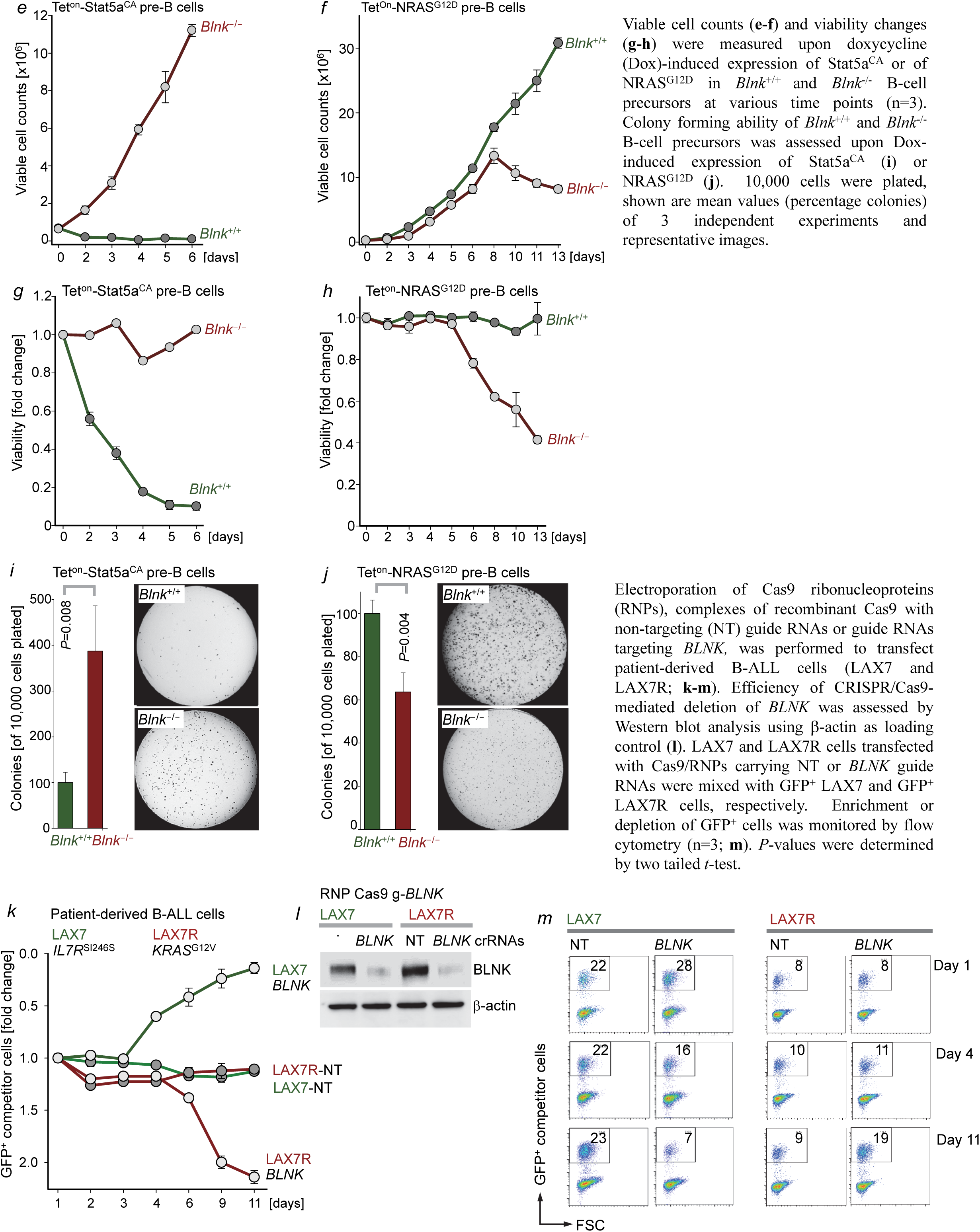
BLNK enables oncogenic ERK-signaling at the expense of STAT5-MYC

**Extended Data Figure 10:**
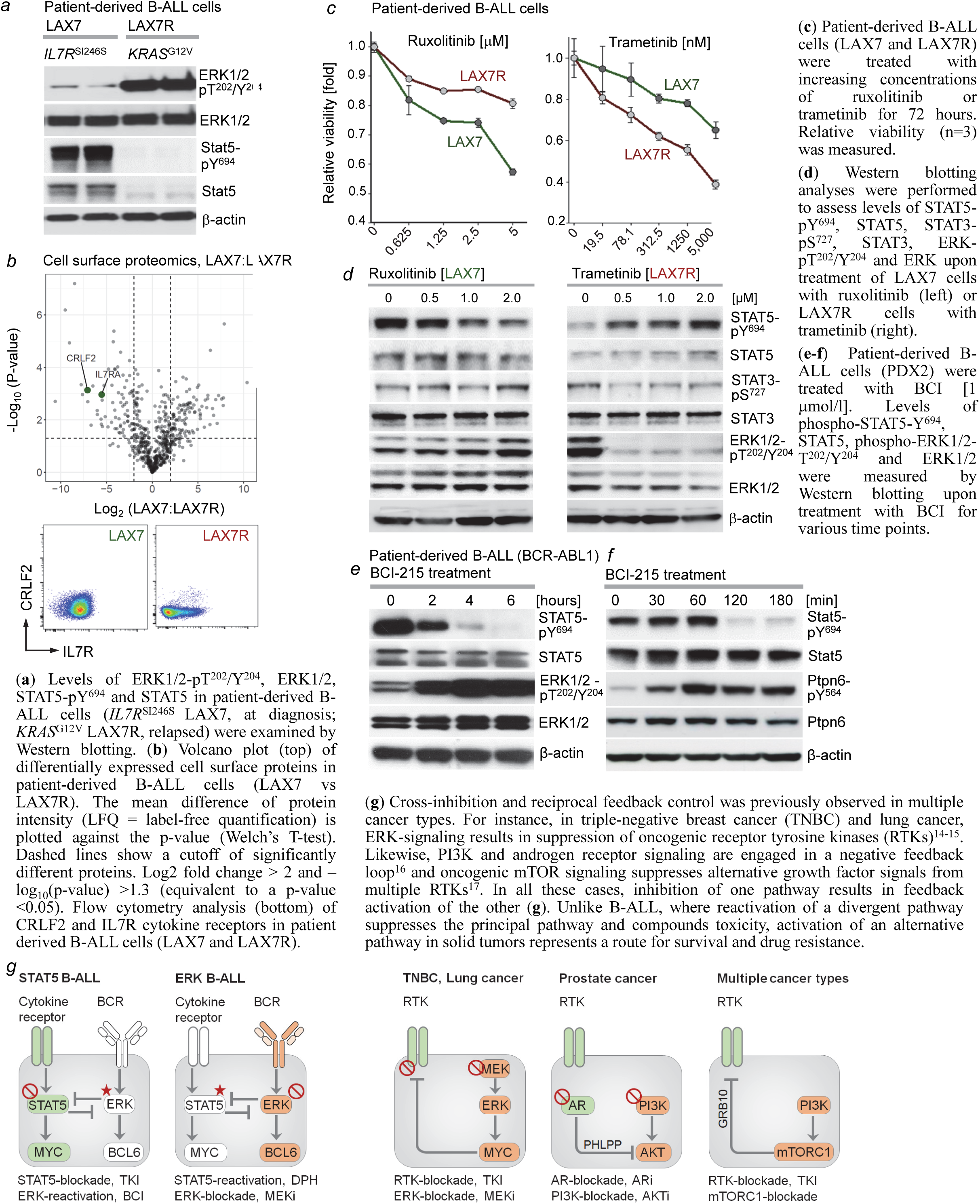
Divergent drug-responses in a STAT5- and ERK-driven pair of primary and relapse B-ALL

## References

1. Fearon, ER et al. Clonal analysis of human colorectal tumors. Science 238, 193–197 (1987)

2. Goetz, C et al. STAT5 activation underlies IL7 receptor-dependent B cell development. J. Immunol. 172, 4770–4778 (2004)

3. Malin, S. et al. Role of STAT5 in controlling cell survival and immunoglobulin gene recombination during pro-B cell development. Nat. Immunol. 11, 171–179 (2010)

4. Russell, LJ et al. Deregulated expression of cytokine receptor gene, CRLF2, is involved in lymphoid transformation in B-cell precursor acute lymphoblastic leukemia. Blood 114, 2688–2698 (2009)

5. Meydan, N et al. Inhibition of acute lymphoblastic leukemia by a Jak2 inhibitor. Nature 379, 645–648 (1996)

6. Hoelbl, A. et al. Stat5 is indispensable for the maintenance of bcr/abl-positive leukaemia. EMBO Mol. Med. 2, 98–110 (2010)

7. Katerndahl, CDS et al. Antagonism of B cell enhancer networks by STAT5 drives leukemia and poor patient survival. Nat. Immunol. 18, 694–704 (2017)

8. Shaw, AC et al. Activated Ras signals developmental progression of recombinase-activating gene (RAG)-deficient pro-B lymphocytes. J. Exp. Med. 189, 123–129 (1999)

9. Irving, J et al. Ras pathway mutations are prevalent in relapsed childhood acute lymphoblastic leukemia and confer sensitivity to MEK inhibition. Blood 124, 3420–3430 (2014)

10. Tiacci, E et al. BRAF mutations in hairy-cell leukemia, N. Engl. J. Med. 364, 2305–2315 (2011)

11. Anderson, LJ et al. EBV LMP2A provides a surrogate pre-B cell receptor signal through constitutive activation of the ERK/MAPK pathway. J. Gen. Virol. 89, 1563–1568 (2008)

12. Feldhahn, N et al. Mimicry of a constitutively active pre-B cell receptor in acute lymphoblastic leukemia cells. J. Exp. Med. 201, 1837–1852 (2005)

13. Rowland, SL et al. Ras activation of Erk restores impaired tonic BCR signaling and rescues immature B cell differentiation. J. Exp. Med. 207, 607–621 (2010)

14. Duncan, JS et al. Dynamic reprogramming of the kinome in response to targeted MEK inhibition in triple-negative breast cancer. Cell 149, 307–321 (2012)

15. Manchado, E et al. A combinatorial strategy for KRAS-mutant lung cancer. Nature 534, 647–651 (2016)

16. Carver, BS et al. Reciprocal feedback regulation of PI3K and androgen receptor signaling in PTEN-deficient prostate cancer. Cancer Cell 19, 575–586 (2011)

17. Hsu, PP et al. The mTOR-regulated phosphoproteome reveals a mechanism of mTORC1-mediated inhibition of growth factor signaling. Science 332, 1317–1322 (2011)

18. Yasuda, T et al. Erk kinases link pre-B cell receptor signaling to transcriptional events required for early B cell expansion. Immunity 28, 499–508 (2008)

19. Shojaee, S et al. PTEN opposes negative selection and enables oncogenic transformation of pre-B cells. Nat. Med. 22, 379–387 (2016)

20. Yang, Y et al. Exploiting synthetic lethality for the therapy of ABC diffuse large B cell lymphoma. Cancer Cell 21, 723–737 (2012)

21. Nikolaev, SI et al. Frequent cases of RAS-mutated Down syndrome acute lymphoblastic leukaemia lack JAK2 mutations. Nat. Commun. 5, 4654 (2014)

22. Herold, T. et al. Adults with Philadelphia chromosome-like acute lymphoblastic leukemia frequently have IGH-CRLF2 and JAK2 mutations, persistence of minimal residual disease and poor prognosis. Haematologica 102, 130–138 (2017)

23. Zhang J. et al. Key pathways are frequently mutated in high-risk childhood acute lymphoblastic leukemia: a report from the Children’s Oncology Group. Blood 118, 3080–3087 (2011)

24. Jerchel, I.S. et al. RAS pathway mutations as a predictive biomarker for treatment adaptation in pediatric B-cell precursor acute lymphoblastic leukemia. Leukemia 32, 931–940 (2018)

25. Harvey R.C. et al. Identification of novel cluster groups in pediatric high-risk B-precursor acute lymphoblastic leukemia with gene expression profiling: correlation with genome-wide DNA copy number alterations, clinical characteristics, and outcome. Blood 116, 4874–4884 (2010).

26. Schwartzman, O et al. Suppressors and activators of JAK-STAT signaling at diagnosis and relapse of acute lymphoblastic leukemia in Down syndrome. Proc. Natl. Acad. Sci. USA 114, E4030–E4039 (2017)

27. Heltemes-Harris, LM et al. Sleeping Beauty transposon screen identifies signaling modules that cooperate with STAT5 activation to induce B-cell acute lymphoblastic leukemia. Oncogene 35, 3454–3464 (2016)

28. Porpaczy, E et al. Aggressive B-cell lymphomas in patients with myelofibrosis receiving JAK1/2 inhibitor therapy. Blood 132, 694–706 (2018)

29. Mandal, M et al. Ras orchestrates exit from the cell cycle and light-chain recombination during early B cell development. Nat. Immunol. 10, 1110–1117 (2009)

30. Duy, C et al. BCL6 is critical for the development of a diverse primary B cell repertoire. J. Exp. Med. 207, 1209–1221 (2010)

31. Swaminathan, S et al. Mechanisms of clonal evolution in childhood acute lymphoblastic leukemia. Nat. Immunol. 16, 766–774 (2015)

32. Walker, S.R. et al. STAT5 outcompetes STAT3 to regulate the expression of the oncogenic transcriptional modulator BCL6. Mol Cell Biol. 33, 2879–2890 (2013)

33. Cerchietti, L.C. et al. A peptomimetic inhibitor of BCL6 with potent antilymphoma effects in vitro and in vivo. Blood 113, 3397–3405 (2009)

34. Cardenas, M.G. et al. Rationally designed BCL6 inhibitors target activated B cell diffuse large B cell lymphoma. J Clin. Invest. 126, 3351–3362 (2016)

35. Korotchenko, VN et al. In vivo structure-activity relationship studies support allosteric targeting of a dual specificity phosphatase. Chembiochem. 15, 1436–1445 (2014)

36. Shojaee, S et al. Erk negative feedback control enables pre-B cell transformation and represents a therapeutic target in acute lymphoblastic leukemia. Cancer Cell 28, 114–128 (2015)

37. Xiao, W et al. Tumor suppression by phospholipase C-beta3 via SHP-1-mediated dephosphorylation of Stat5. Cancer Cell 16, 161–171 (2009)

38. Fusaki, N et al. BLNK is associated with the CD72/SHP-1/Grb2 complex in the WEHI231 cell line after membrane IgM cross-linking. Eur. J. Immunol. 30, 1326–1330 (2000)

39. Imamura, Y et al. BLNK binds active H-Ras to promote B cell receptor-mediated capping and ERK activation. J. Biol. Chem. 284, 9804–9813 (2009)

40. Druker, BJ et al. Effects of a selective inhibitor of the Abl tyrosine kinase on the growth of Bcr-Abl positive cells. Nat. Med. 2, 561–566 (1996)

41. Das Thakur, M et al. Modelling vemurafenib resistance in melanoma reveals a strategy to forestall drug resistance. Nature 494, 251–255 (2013)

42. Yang, J et al. Discovery and characterization of a cell-permeable, small-molecule c-Abl kinase activator that binds to the myristoyl binding site. Chem. Biol. 18, 177–186 (2011)

43. Emery, CM et al. MEK1 mutations confer resistance to MEK and B-RAF inhibition. Proc. Natl. Acad. Sci. USA 106, 20411–20416 (2009)

44. Hanahan, D et al. The hallmarks of cancer. Cell 100, 57–70 (2000)

45. Müschen, M. Autoimmunity checkpoints as therapeutic targets in B cell malignancies. Nat. Rev. Cancer. 18, 103–116 (2018)

